# RBM10 loss induces aberrant splicing of cytoskeletal and extracellular matrix mRNAs and promotes metastatic fitness

**DOI:** 10.1101/2024.07.09.602730

**Authors:** Gnana P. Krishnamoorthy, Anthony R. Glover, Brian R. Untch, Nickole Sigcha-Coello, Bin Xu, Dina Vukel, Yi Liu, Vera Tiedje, Katherine Berman, Prasanna P. Tamarapu, Adrian Acuña-Ruiz, Mahesh Saqcena, Elisa de Stanchina, Laura Boucai, Ronald A. Ghossein, Jeffrey A. Knauf, Omar Abdel-Wahab, Robert K. Bradley, James A. Fagin

## Abstract

RBM10 modulates transcriptome-wide cassette exon splicing. Loss-of-function *RBM10* mutations are enriched in thyroid cancers with distant metastases. Analysis of transcriptomes and genes mis-spliced by RBM10 loss showed pro-migratory and RHO/RAC signaling signatures. RBM10 loss increases cell velocity. Cytoskeletal and ECM transcripts subject to exon-inclusion events included vinculin (VCL), tenascin C (TNC) and CD44. Knockdown of the VCL exon inclusion transcript in *RBM10*-null cells reduced cell velocity, whereas knockdown of TNC and CD44 exon-inclusion isoforms reduced invasiveness. RAC1-GTP levels were increased in *RBM10*-null cells. Mouse *Hras^G12V^/Rbm1O^KO^* thyrocytes develop metastases that are reversed by RBM10 or by combined knockdown of VCL, CD44 and TNC inclusion isoforms. Thus, *RBM10* loss generates exon inclusions in transcripts regulating ECM-cytoskeletal interactions, leading to RAC1 activation and metastatic competency. Moreover, a CRISPR-Cas9 screen for synthetic lethality with RBM10 loss identified NFkB effectors as central to viability, providing a therapeutic target for these lethal thyroid cancers.

**SUMMARY:** RNA splicing factor mutations are common in cancer but connecting phenotypes to specific misspliced genes has been challenging. We show that RBM10 loss leads to exon inclusions of transcripts regulating ECM-cytoskeletal interactions and RAC1-GTP activation, sufficient to promote metastatic fitness.

## INTRODUCTION

Most multi-exon human genes undergo alternative splicing (AS), which vastly expands the repertoire of the human proteome (Reixachs-Sole and Eyras, 2022). AS yields multiple isoforms from a single pre-mRNA molecule by exon skipping (or cassette exon exclusion), mutually exclusive exons, alternative 5’ donor sites, alternative 3’ acceptor sites, intron retention and intron splicing (Lopez, 1998, Croft et al., 2000). Although AS is a common mechanism of gene regulation, the determinants of lineage-specific transcript diversity and how cells maintain stoichiometric control over various isoforms is incompletely understood both in normal and tumorigenic states (Bradley and Anczuków, 2023). Cancer genome studies identified recurrent loss-of-function and hotspot mutations in splicing genes including *SF3B1*, *U2AF1*, *SRSF2*, *FUBP1*, and *RBM10* (Seiler et al., 2018). These lesions exert transcriptome-wide deregulation of mRNA splicing that promote tumorigenesis through complex mechanisms that are difficult to unravel, since they impact on multiple gene targets.

Germline alterations of *RBM10* in humans results in TARP syndrome in males (OMIM #311900) characterized by **t**alipes equinovarus (club foot), **a**trial septal defect, **r**obin sequence (micrognathia), and **p**ersistent left superior vena cava. Other manifestations include craniofacial abnormalities, hypotonia and developmental delay. The underlying mechanism of disease is not well defined, but embryonic expression of RBM10 in the mouse is found in the branchial arches and limb buds (Johnston et al., 2014), sites where regulation of cell motility play a fundamental role. The *RBM10* gene, a key player in AS that is not a component of the core spliceosome, is located on the X-chromosome and is mutated in 10% of NSCLC, 8% of bladder cancers and 11% of non-anaplastic thyroid cancers who died of metastatic disease (Ibrahimpasic et al., 2017, Seiler et al., 2018). Moreover, MSK-MET, an integrated pan-cancer cohort study of tumor genomic and clinical outcome data of over 25,000 patients, showed that *RBM10* alterations associate with metastatic burden in papillary thyroid cancer (PTC) but not in other tumor types (Nguyen et al., 2022). The specific AS targets of RBM10 that may mediate this metastatic phenotype are unknown.

CLIP-seq and RNASeq have been used to identify the splicing targets of RBM family proteins, including RBM10, primarily in HEK293 and HeLa cells (Wang et al., 2013, Bechara et al., 2013). RBM10 binds to splice sites of specific exons and promotes cassette exon exclusion from target pre-mRNAs, and to a lesser extent other AS events (Wang et al., 2013, Inoue et al., 2014). *RBM10* mutant tumors have globally reduced mRNA levels compared to normal (Seiler et al., 2018). Bladder and non-small cell lung cancer (NSCLC) with *RBM10* mutations are enriched for cassette exon inclusions, which when in frame can alter the function of a protein. Some of these have been implicated in pathogenesis of specific cancer types. Upon RBM10 loss an exon inclusion event in NUMB mRNA leads to decreased NUMB protein levels through hyper-ubiquitination and destabilization and consequent de-repression of NOTCH signaling in NSCLC cells (Bechara et al., 2013). In addition, an exon 5 inclusion isoform of EIF4H, which encodes a eukaryotic translation initiation factor, has been shown to be a target of *RBM10* loss in lung adenocarcinoma tissues and to mediate in part the growth inhibitory effects of RBM10 overexpression in NSCLC lines (Zhang et al., 2020). A number of other genes including *CREBBP (Wang et al., 2013), BID (Bechara et al., 2013), FAS, BCL-XL (Inoue et al., 2014)* and *SMN2 (Sutherland et al., 2017)* have been shown to be AS targets of RBM10 and implicated in tumorigenesis, primarily based on *in vitro* silencing of RBM10 in cancer cell lines or HeLa cells (Zhao et al., 2017), (Inoue et al., 2014).

In this study we demonstrate the molecular and biological consequences of *RBM10* loss in thyroid cancer employing human cell lines with naturally occurring *RBM10* lesions and in cells derived from a GEMM of metastatic thyroid cancer driven by *Rbm10* loss. We find that *RBM10* loss leads to AS of pre-mRNAs that underpin development of metastases. We reveal a link between *RBM10* loss and expression of a specific set of cytoskeletal and extracellular matrix (ECM) isoforms that converge to induce cell motility, invasiveness and metastatic fitness, primarily through activation of the RAC1 signaling pathway. Additionally, a genome-wide CRISPR-Cas9 screen in RBM10-mutant KTC1 cells identified dropout genes in the NFκB signaling pathway, revealing a sensitivity to NFκB small molecule inhibitors.

## RESULTS

### Prevalence of *RBM10* mutations in thyroid cancer

Papillary thyroid cancers (PTC) are mostly indolent well-differentiated tumors associated with good clinical outcomes. Of the 496 patient tumors genotyped in TCGA study of PTC, none had *RBM10* mutations (Fig. 1A). Of note, in that series only 8 patients had distant metastases at presentation. By contrast the prevalence of *RBM10* mutations in PTC from the MSK clinical genomic database (MSK-IMPACT), which is enriched for patients with recurrent metastatic disease, was 3.98% (15/377). In high-grade follicular cell-derived thyroid cancer (HGFCTC) the prevalence was 6.7% (15/224) and in anaplastic thyroid cancers (ATC) 1.47% (2/136) (Fig. 1A). In the combined TCGA and MSK-clinical cohorts *RBM10* alterations are significantly enriched (30/553; p<0.0001) in non-anaplastic thyroid cancer patients (PTC and HGFCTC) with distant metastatic disease as compared to those without distant metastases (0/514) (Fig. 1B). Most *RBM10* alterations in thyroid cancer are frameshift or non-sense mutations (Fig. S1A), consistent with its role as a tumor suppressor. *RBM10* loss co-occurs with MAPK pathway driver oncoproteins, most commonly with *RAS* (43%) and *TERT* promoter mutations (81%) and shows a trend towards mutual exclusivity with *TP53* mutations (Fig. S1B-C).

**Figure 1:**
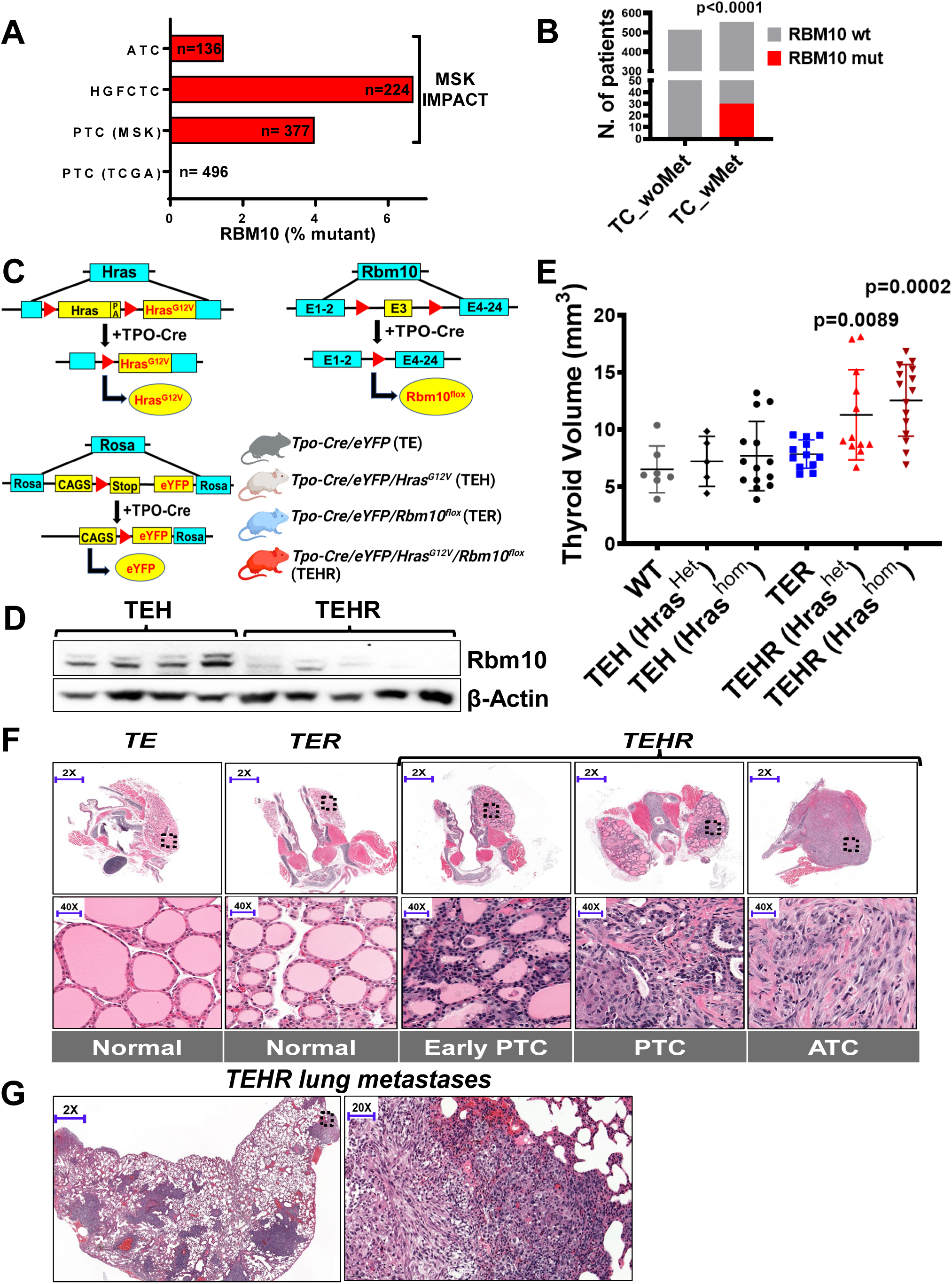
*Thyroid cancer phenotypes associated with RBM10 mutations:* (A) Frequency of *RBM10* mutations in histological subtypes of thyroid cancer samples from MSK-clinical cohort (n=737) sequenced by MSK-IMPACT and PTC from TCGA (n=496). PTC: papillary thyroid cancer; HGFCTC: High grade follicular cell-derived thyroid cancer; ATC: anaplastic thyroid cancer. (B) Proportion of *RBM10* mutation in non-anaplastic (PTC+HGFCTC) thyroid cancers from MSK-IMPACT and TCGA showing association of *RBM10* mutation in patients with distant metastases; 2-sided Fisher’s exact t-test p<0.0001. (C) Diagram of *Hras, Rbm10 and EYFP* transgenes in the mouse model. In the presence of *Tpo-Cre*, YFP and mutant Hras are expressed and Rbm10 is inactivated in thyrocytes. (D) Western blot showing Rbm10 expression in mouse thyroid lysates from the indicated genotypes. (E) Thyroid volume measurement by ultrasound of 25-week-old mice for the indicated genotypes; p values derived by unpaired t test vs WT. (F) Low (top) and high power (bottom) magnification of H&E-stained thyroids of the indicated genotypes showing representative histology. (G) H&E-stained sections of thyroid cancer metastases to lung from a *TEHR* mouse; low (left) and high magnification (right).

### *Rbm10* loss promotes mouse thyroid cancer development in the context of oncogenic *Hras*

To investigate the role of *Rbm10* loss in thyroid cancer biology we developed compound mice with thyroid-specific knockout of *Rbm10* either alone or in the context of a knock-in allele of Hras^G12V^ (Fig. 1C). The *Rbm10* floxed allele (*Rbm10^flox^*) results in a non-functional transcript following Cre recombinase-mediated excision of exon 3 (Wang et al., 2023). We crossed *Rbm10^flox^* mice with *Tpo-Cre/Caggs-LSL-eYFP* (“TE”) to generate *Tpo-Cre/Caggs-LSL-eYFP/ Rbm10^flox^* (“TER”) mice. We then crossed TER with *Tpo-Cre/ Caggs-LSL-eYFP /FR-Hras^G12V^*(“TEH”) mice, which harbor floxed tandem *Hras* alleles that upon recombination generate mice with endogenous expression of *Hras^G12V^* in thyroid follicular cells (Chen et al., 2009), to generate *Tpo-Cre/Caggs-LSL-eYFP//FR-Hras^G12V^/ Rbm10^flox^* (“TEHR”) mice (Fig. 1C). TEHR mice result in thyroid-specific loss of Rbm10 (Fig. 1D) and expression of Hras^G12V^ and YFP. TER and TEH mice showed no increase in thyroid volume or thyroid cancer development through 12 months of life (Fig. 1E and Fig. S1D). As previously reported, a subset of TEH mice developed mild thyroid hyperplasia after a prolonged latency (Montero-Conde et al., 2017). By contrast, TEHR mice developed a spectrum of thyroid cancer phenotypes by 10-12 months (Fig. 1F and Fig. S1D): 9% had infiltrative tumors without well-developed features of any subtype of human differentiated thyroid cancer, which we termed “early PTC” based on their nuclear features; 12% developed frank PTC and 76% were ATC-like cancers characterized by pleomorphic nuclei and spindle cell metaplasia. In addition, 18% of this cohort had lung metastases (Fig. 1G and Fig. S1D).

### *RBM10* loss induces cassette exon inclusion isoforms of ECM/cytoskeletal remodeling genes

To study the molecular processes underpinning tumorigenic events driven by *RBM10* mutations we generated 5 isogenic human thyroid cancer cell lines. Two of these are *RBM10*- null with doxycycline-induced expression of RBM10: KTC1, derived from a male with two somatic *RBM10* mutations in the same allele (RBM10_p.A577fs and p.S580R) and PE121410, from a female with an RBM10_p.Q344fs mutation. Hth83, SW1736, 8505C are *RBM10* wild type cell lines with shRNA-mediated silencing of RBM10 (Fig. 2A). Four of the 5 cell lines showed an increase in cell proliferation in RBM10-null or knockdown (KD) compared to their respective isogenic RBM10-expressing controls (Fig. S2A), suggesting that RBM10 loss confers a growth advantage to the cells. The effects of RBM10 loss on growth and metastatic propensity may result from common AS targets driving both mechanisms of tumorigenesis or from distinct sets of AS events. We took a two-pronged approach to investigate this: 1. High depth RNAseq to uncover AS targets of RBM10 loss potentially involved in promoting metastases; 2. A CRISPR/Cas-9 genome-wide depletion screen designed to identify genes involved in cell growth in RBM10 deficient cells (Fig. 2B). AS analysis of the RNAseq using the MISO splicing analysis model (Katz et al., 2010) identified 113 alternatively spliced RBM10 targeted genes common to the 5 isogenic cell lines (Fig. S2B). The most common AS events associated with RBM10 loss were exon inclusions (SE) (Fig. 2C), followed by alternate usage of 3’splice sites (A3SS), alternate usage of 5’splice sites (A5SS) and retained introns (RI). The predilection for alternative usage of 3’ splice sites (A3SS) is consistent with the preferential binding of RBM10 to the vicinity of 3’ intron-exon boundaries (Wang et al., 2013). In addition, we employed Partek Flow Alt-Splicing Analysis, which identified 1005 genes with 1456 differentially spliced isoforms (Table S1) common to PE121410 and KTC1 cells, including 79/112 of those identified by the MISO approach. We consider these two cell lines as the most informative because they had naturally occurring *RBM10* mutations. Gene Ontology (GO) analysis confined to genes subject to AS by *RBM10* loss show enriched GO terms linked to ECM/cytoskeletal remodeling process, with the top GO terms being regulation of focal adhesion assembly, positive regulation of actin filament bundle assembly and positive regulation of cytoskeleton organization (Fig. 2D). In addition to the previously reported genes containing cassette-exon splicing targets of RBM10, such as NUMB (Hernandez et al., 2016), SMN2 (Sutherland et al., 2017) and EIF4H (Zhang et al., 2020), ECM/cytoskeletal modulating genes were among the top differentially spliced transcripts: e.g. VCL, CD44, FN1, TPM1 and TPM3 (Fig. 2E).

**Figure 2:**
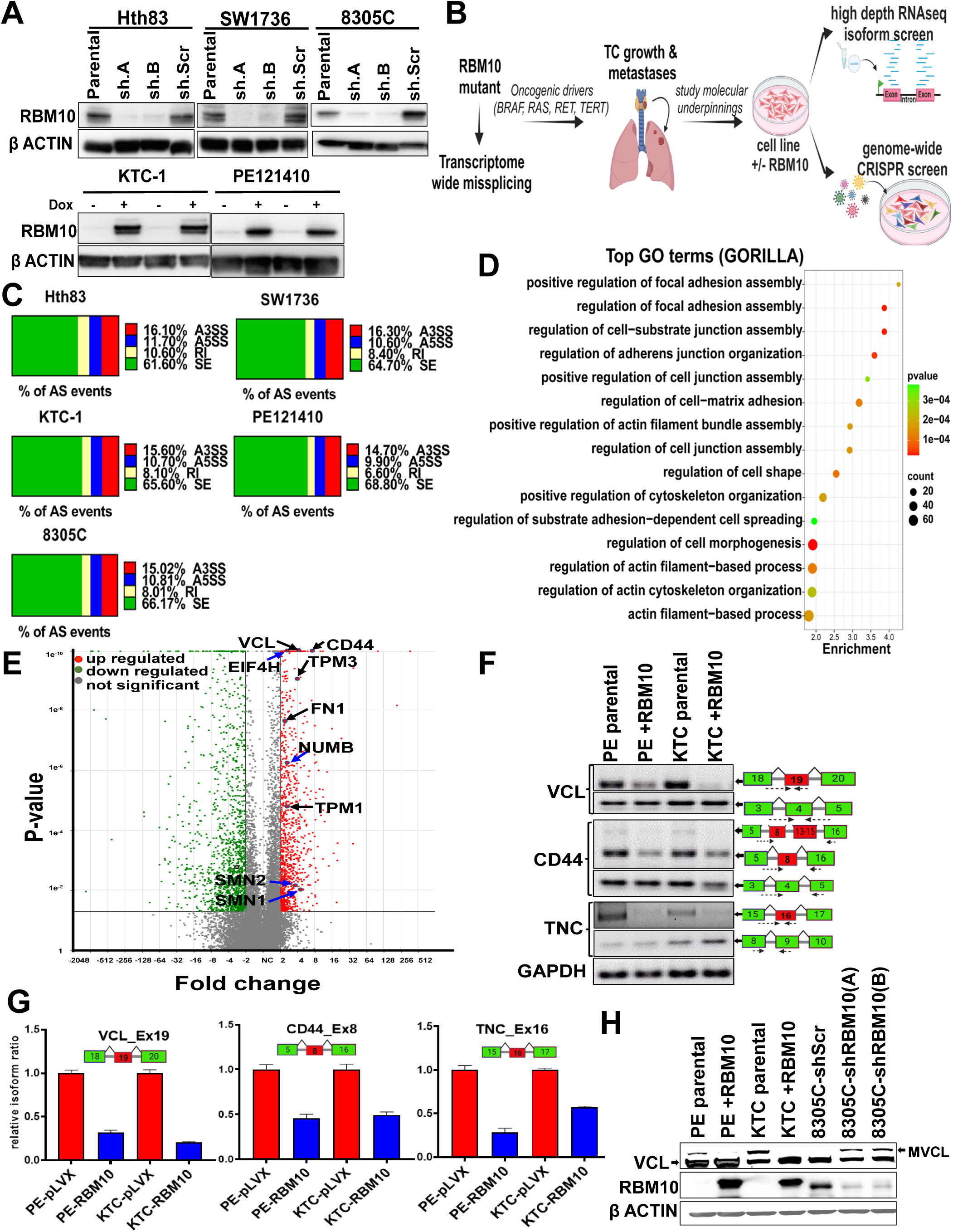
*RBM10 regulates AS of ECM and cytoskeletal transcripts:* (A) Western blot of RBM10 in isogenic human thyroid cancer cell lines. (B) Experimental strategy to identify molecular mechanisms of thyroid cancer growth and metastases in RBM10-deficient cells; Illustration by Biorender. (C) Average changes in aberrant splicing (AS) events in RBM10 deficient detected by high depth RNAseq in the RBM10 isogenic cell lines; SE – cassette exon inclusion isoforms; RI – retained intron; A3SS and A5SS – most intron-proximal isoform for competing 3’ or 5’ splice sites. (D) Gene ontology using ranked differentially spliced genes by GORILLA showing enrichment in pathways involved in cell adhesion and cytoskeleton reorganization. (E) Volcano plot showing differentially expressed alternative spliced isoforms detected by Partek flow RNAseq alt-splicing analysis in the combined isogenic PE121410 and KTC1 cells. Arrows point to representative isoforms of the indicated genes. Black arrows: ECM/cytoskeletal modifying transcripts. Blue arrows: known RBM10 AS genes. (F) RT-PCR showing indicated exon inclusion isoforms of VCL, CD44 and TNC in PE121410 and KTC1 cells and their decrease with RBM10 expression. Constitutive isoforms and GAPDH were used as controls. Red - cassette exon; Green - constitutive exon; Dotted arrow - primer placement. (G) Relative ratio of the indicated inclusion isoforms of VCL, CD44 and TNC determined by qRT-PCR in PE121410 and KTC1 cells -/+ RBM10. (H) Western blot of vinculin and β actin in the indicated cells. Arrow points to VCL and MVCL.

We identified the RBM10-regulated cassette exons based on the RNAseq data (Fig. S2C-D) and validated the inclusion isoform transcripts by qRT-PCR using junction-specific primers, normalized to the respective gene-specific expression using primers against constitutive exons (Fig. 2F-G). This revealed the following ECM/cytoskeletal genes with cassette exons regulated by RBM10: 1. Exon 19 inclusion of VCL (hereafter called as VCL_Ex19) (Fig. 2H); 2. Exon 8 inclusion isoforms of CD44 (collectively called as CD44_Ex8), one consisting of the exon 8 inclusion alone and the other with combined inclusions of exon 8 and exons 13-15 (Fig. 2F-G); 3. Exon16 inclusion isoform of TNC (TNC_Ex16) (Fig. 2F-G) .

Vinculin (VCL) is a modulator of focal adhesion assembly, anchorage of F-actin to the membrane and of cytoskeletal organization (Lee et al., 2019). VCL is encoded by 21 exons (Fig. S2E). Inclusion of the cassette exon 19 in RBM10-null cells generates an isoform called meta- vinculin (MVCL or VCL_Ex19 in this manuscript), characterized by a 68-residue insertion in the actin-binding tail of the protein (Byrne et al., 1992). MVCL is expressed in cardiac and smooth muscle at sub-stoichiometric levels relative to VCL. MVCL expression leads to fewer but larger focal adhesions per cell and to enhancement of cell motility (Lee et al., 2019). Germline mutations leading to substitutions within the MVCL 68-aminoacid insertion are associated with congenital cardiomyopathies (Olson et al., 2002). Our data point to a central role for RBM10 in regulating the alternative splicing of VCL mRNA, thus controlling the balance of VCL to MVCL expression levels (Fig. 2H).

CD44 is a multifunctional cell surface adhesion non-kinase receptor involved in cell-cell and cell-matrix interactions, encoded by 20 exons, 10 of which are constitutive (first 5 and last 5) and 10 that are variable (middle 10) (NaorSionov and Ish-Shalom, 1997) (Fig. S2E) . The encoded standard (CD44_s_) and variable (CD44_v_) isoforms are implicated in the biology of multiple cancer types (Chen et al., 2018). We show that two of the exon-8 inclusion isoforms (CD44_v3,_ and CD44_v3_, _v8-10_; collectively called as CD44_EX8 in this manuscript) are AS targets of RBM10 (Fig. 2F and Fig. S2F).

Tenascin C (TNC), one of the candidates from the MISO analysis, is encoded by 28 exons, 7 of which are variable (Fig. S2E). RT-PCR quantification of each variable exon revealed that *RBM10*-null cells differentially expressed a transcript including exon 16 (TNC_EX16) (Fig. 2F and Fig. S2F), which encodes a tandemly arrayed fibronectin (FN) binding domain that promotes cell detachment by competing with FN for interaction with integrins (Huang et al., 2001).

### *RBM10* loss promotes metastatic properties mediated by exon inclusion isoforms of VCL, CD44 and TNC

To explore potential mechanisms underpinning the effects of RBM10 loss on thyroid cancer metastases we performed Ingenuity Pathway Analysis (IPA) on bulk RNA sequencing on parental and RBM10-expressing isogenic PE121410 cells. This revealed a strong signal for functional annotations involving cell migration and invasion (Fig. 3A) and for modulation of ECM and cytoskeleton effector pathways: e.g. actin nucleation, actin-based motility, remodeling of epithelial adherens junctions, integrin-, actin cytoskeletal- and Rho GTPase-signaling (Fig. 3B). Prompted by these findings we next asked whether RBM10 loss conferred cells with a pro-migratory phenotype. For this we monitored individual cell migration tracks employing time lapse imaging and calculated migration velocity in the RBM10 isogenic KTC1, PE121410 and 8305C cell lines (Fig. 3C). Cells with endogenous *RBM10* loss-of-function mutations (KTC1, PE121410) and shRNA mediated KD of RBM10 display longer cell migratory tracks (Fig. 3D) compared to the respective RBM10 expressing comparators, and higher mean migration velocity (Fig. 3E).

**Figure 3:**
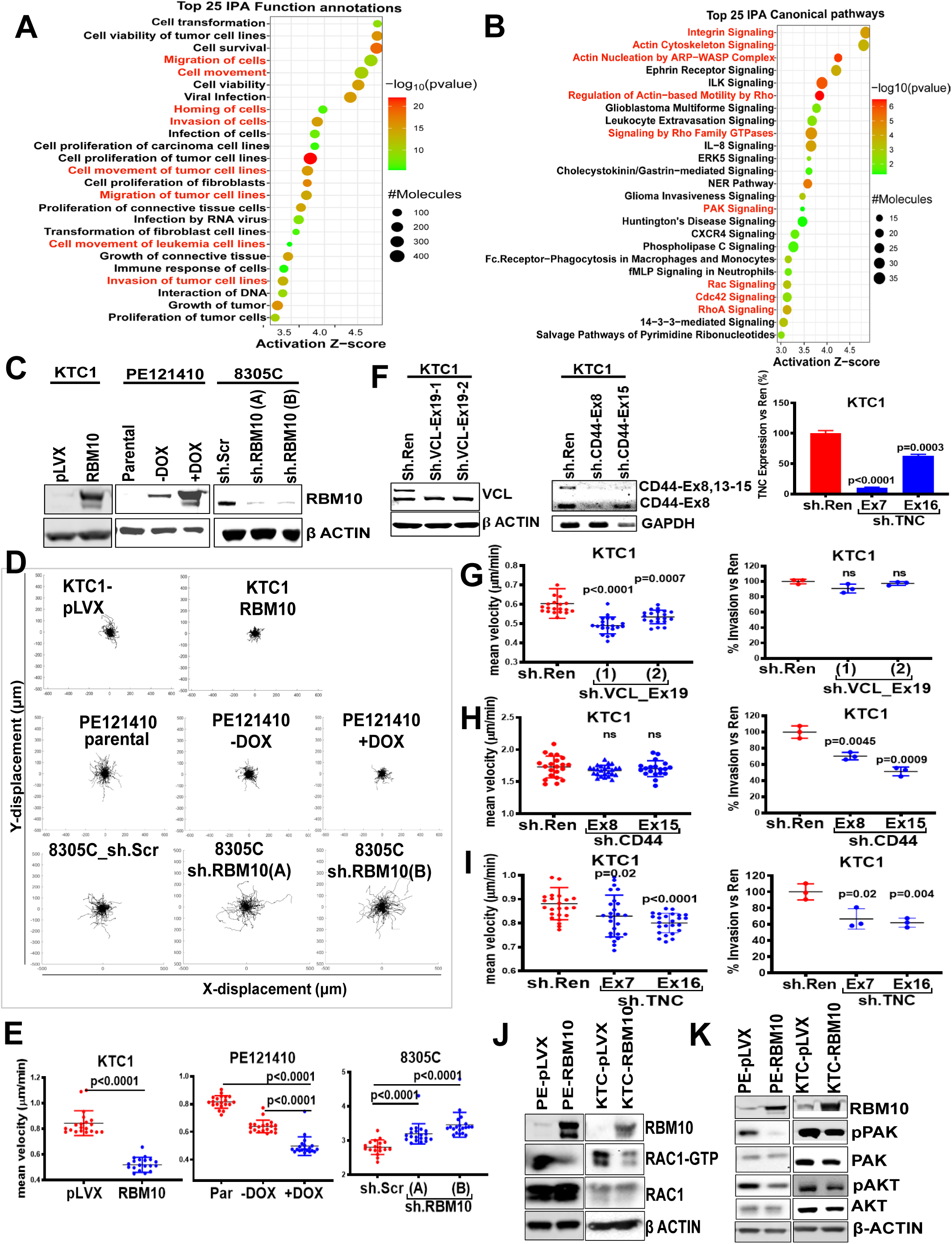
*Impact of RBM10-induced AS of VCL, CD44 and TNC on cell motility and invasiveness:* IPA of RNAseq of RBM10-null vs RBM10-expressing PE121410 cells showing top 25 functional annotations (A) and canonical pathways (B). Highlighted in red are pathways involved in cell migration, movement, invasiveness, RHO-RAC signaling and ECM-cytoskeleton interactions. (C) Western blotting for RBM10 and β Actin in the indicated cell lines. (D) Top two rows: Representative cell trajectories by time lapse imaging of RBM10-null KTC1 and PE121410 at baseline and after dox-induction of RBM10 for 48h. Third row: Cell trajectory of 8305C cells transduced with scrambled or RBM10 shRNAs. (E) Mean cell velocity quantified by time lapse imaging for the 3 cell lines -/+ RBM10. (F) Western blot (*left*), RT-PCR (*middle*) and qRT-PCR (*right*) for VCL, CD44 and TNC, respectively, in KTC1 cells transduced with sh.Renilla or sh.RNAs targeting the indicated exons demonstrating isoform-specific KDs. (G-I) Cell migration velocity by time lapse imaging (*left*) and cell invasion by transwell assay (*right*) in control vs. isoform-specific KD of VCL (G), CD44 (H) and TNC (I); p values by unpaired t test vs sh.Ren. (J & K) Increased RAC1-GTP levels (J) and RAC1 downstream signaling (pPAK1, pAKT-S473) (K) in RBM10-null PE121410 and KTC1 cells.

We next asked whether individual ECM and cytoskeletal exon-inclusion events driven by RBM10 loss impacted cell migration and/or invasion. For this we created isoform-specific stable KD with short hairpins against the inclusion exons of VCL (sh.VCL_Ex19), CD44 (sh.CD44_Ex8, sh.CD44_Ex11 and sh.CD44_Ex15) and TNC (sh.TNC_Ex7, sh.TNC_Ex16) in KTC1 cells (Fig. 3E-G). Knockdown of the Ex19 inclusion isoform of vinculin with two distinct shRNAs silenced expression of metavinculin and led to decreased cell migration velocity but elicited no effect on invasiveness as assessed by time lapse imaging and a transwell invasion assay, respectively (Fig. 3F-G). By contrast, KD of CD44_Ex8 inclusion isoforms decreased cell invasion without impacting migration velocity (Fig. 3F and Fig. 3H), whereas KD of a constitutive TNC exon (TNC_Ex7) as well as the TNC_EX16 inclusion isoform decreased both migration and invasion (Fig. 3F and Fig. 3I). Based on this we hypothesized that RBM10 target genes may contribute to metastatic fitness by illegitimately expressing cassette exon inclusion isoforms of protein intermediates that collectively modulate ECM/cytoskeletal interactions.

### RBM10 loss leads to constitutive activation of RAC1-GTP levels

Rho GTPases relay extracellular signals to regulate actin dynamics, gene transcription, cell cycle progression, cell adhesion, motility and invasion (Jaffe and Hall, 2005, Tang et al., 2008). Among Rho GTPases, RAC1 plays a major role in cell motility by promoting lamellipodia formation, focal adhesions and matrix metalloprotease expression (Ridley et al., 1992). RBM10-null human PE121410 and KTC1 cells transduced with empty vector have markedly elevated RAC1-GTP levels, which are suppressed following RBM10 expression (Fig. 3J). Increased RAC1-GTP levels in PE121410 cells result in phosphorylation of its downstream effectors, including PAK and AKT, which was attenuated by RBM10 expression (Fig. 3K). Interestingly, we previously reported that KTC1 cells harbor an endogenous *RAC1-D63V* mutation (Landa et al., 2016). The D63 residue lies within the highly conserved switch II region of RAC1 that is involved in effector binding to PAK, WASP, and ACK (Mott et al., 1999). Based on the RAC1 crystal structure, D63 is also in direct contact with the binding site for RAC1-GAP (Stebbins and Galan, 2000). Thus, D63 could contribute to both effector function and regulation of GDP-GTP exchange, and mutated residues at this site would be predicted to be activating in nature (Caye et al., 2015, Murphy et al., 2021). Accordingly, expression of RBM10 in KTC1 cells partially inhibited downstream effectors of RAC1-GTP compared to isogenic PE121410 cells.

### ECM and cytoskeletal splicing targets of *RBM10* govern metastatic propensity

We next tested the role of Rbm10 and of mouse exon inclusion isoforms that were orthologous to their corresponding human genes (i.e. Vcl_Ex19 (MVcl); Cd44_EX8; Tnc_EX14) on development of metastases *in vivo*. For this we used a cell line derived from a lung metastasis of an *TEHR* mouse thyroid cancer (64860M), on which we performed rescue experiments by either Rbm10 re-expression or by knocking down the cassette exon inclusion isoforms, alone or in combination (Fig. 4A-B). Expression of Rbm10 decreased migration velocity (Fig. 4C) and invasiveness (Fig. 4D) compared to empty vector-transfected cells. Accordingly, *in vivo* metastatic efficiency of Luc+ 64860M isogenic cells via either tail vein (TV) or orthotopic injection into mouse thyroid showed that Rbm10 re-expression decreased metastatic efficiency (Fig. 4E-F and Fig. S4A-B). We next investigated whether knockdown of the three exon inclusion isoforms, either individually or in combination, impacted metastatic efficiency of 64860M cells. For this we generated stable KDs in Luc+64860M cells of the exon inclusion isoforms individually: shVcl_Ex19, shCd44_Ex8 and shTnc_Ex14; as double KDs: shVcl_Ex19/shCd44_Ex8, shVcl_Ex19/shTnc_Ex14 and shCd44_Ex8/shTnc_Ex14; or as a triple KDs - shVcl_Ex19/shCd44_Ex8/shTnc_Ex14 (Fig. S4C-E). The individual KDs did not show a significant difference in metastatic efficiency *in vivo* as tested by TV injection (Fig. S4F). Short-hairpins against the combined VCL_Ex19/Tnc-Ex14 inclusion isoforms partially suppressed metastases *in vivo*, whereas the other two combinations (sh.Vcl_Ex19/CdD44_Ex8 and sh.Cd44_Ex8/Tnc_Ex14) showed a non-significant inhibitory trend (Fig. 4G). Consistent with this, time lapse imaging showed that the shVcl_Ex19/shTnc_Ex14 double KD was the most effective at decreasing cell motility amongst individual and double KDs tested (Fig. 4H). Notably, the triple KD had the most potent metastases suppressive effect *in vivo* (Fig. 4G).

**Figure 4:**
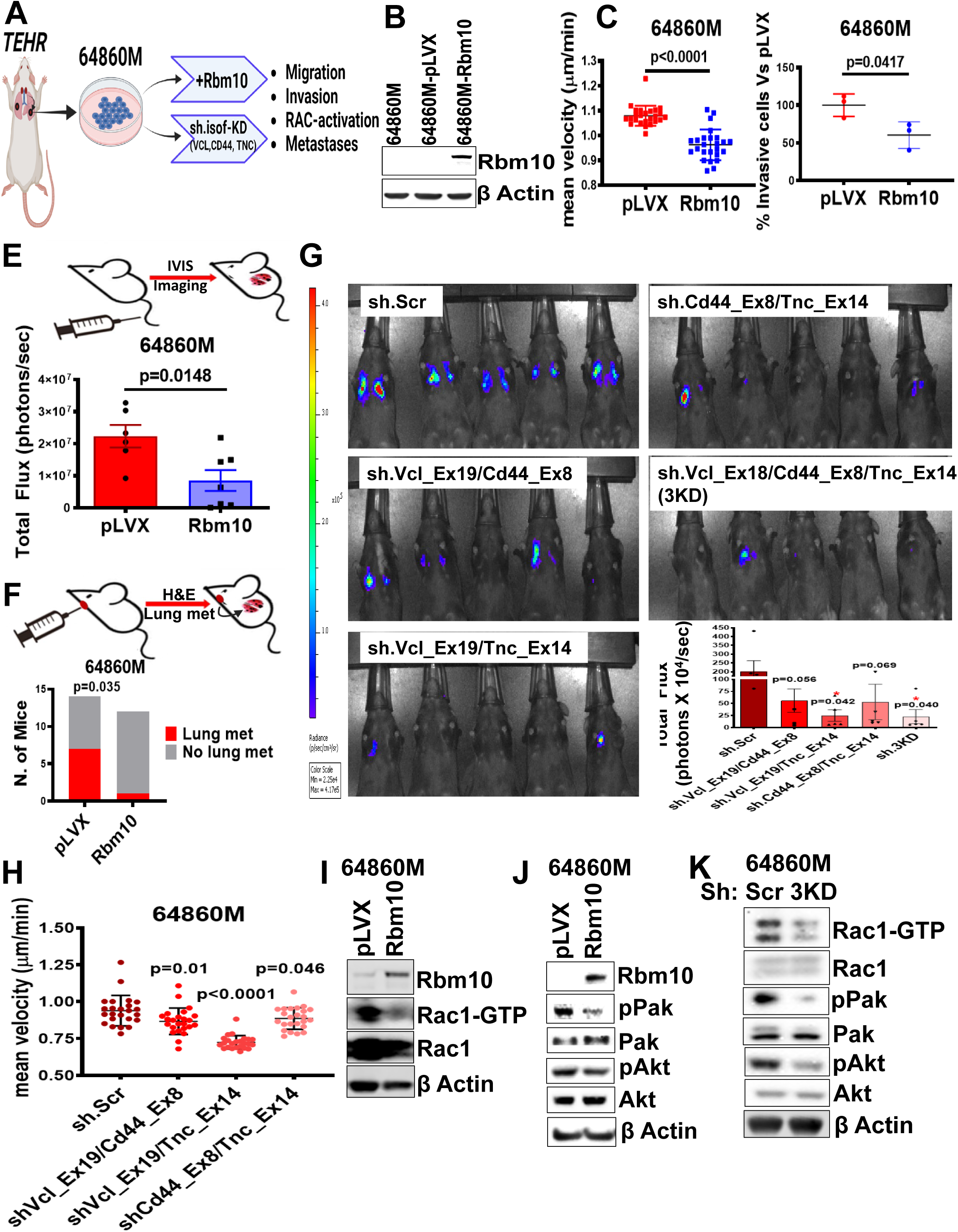
*RBM10-induced exon inclusions of ECM and cytoskeletal transcripts govern metastatic propensity in vivo.* (A) Scheme of *Hras^G12V^/Rbm10^flox^* mouse thyroid cancer lung metastases- derived cell line (64860M) and phenotype rescue experiments by Rbm10 re-expression or isoform- specific KD of Vcl (sh.Vcl_Ex19), Cd44 (sh.Cd44_Ex8) and Tnc (sh.Tnc_Ex14); Illustration by Biorender. (B) Western blot of 64860M cells expressing empty vector (pLVX) and Rbm10. (C) Mean cell migration velocity by time-lapse imaging in 64860M cells -/+ Rbm10. (D) Transwell matrigel invasion assay in 64860M cells -/+ Rbm10. (E) Effect of Rbm10 expression in Luc+ 64860M cells on lung bioluminescence quantification 2 weeks after tail vein injection; p value by unpaired t test. (F) Effect of Rbm10 expression on number of mice with lung metastases assessed by lung H&E, 3-4 weeks after orthotopic implantation of Luc+ 64860M cells into the thyroid; p value by 2-sided Fisher’s exact t-test. (G) Bioluminescence imaging and quantitation of tail vein-injected Luc+ 64860M cells with or without dual and triple isoform-specific KD of Vcl (Vcl_Ex19), Cd44 (Cd44_Ex8) and Tnc (Tnc_Ex14); p value by unpaired t test vs sh.Scr. (H) Vcl, Cd44 and Tnc dual isoform specific KDs in 64860M cells show decreased cell migration velocity compared to sh-Scr transfected cells; p value by unpaired t test vs sh.Scr. (I & J) Decreased Rac1-GTP levels (H) and Rac1 downstream signaling (J) in 64860M cells expressing Rbm10. (K) Isoform- specific triple KD of Vcl, Cd44 and Tnc (sh.3KD) show decreased Rac1-GTP and downstream signaling.

As all three RBM10 target genes studied here operate upstream of Rho-GTPase signaling, we tested the impact of the triple KD on RAC1 activation in 64860M cells. Empty vector-transfected 64860M cells have high Rac1-GTP levels and downstream signaling that is suppressed by dox-induced Rbm10 expression (Fig. 4I-J). Moreover, 64860M-sh.3KD cells had attenuated RAC1- GTP levels and downstream effector signaling compared to a scramble hairpin control (Fig. 4K), providing a plausible mechanistic relationship between ECM and cytoskeletal AS targets of RBM10 loss, RAC1 activation, downstream signaling, cell motility, invasiveness, and metastatic propensity.

### RBM10-loss confers synthetic lethal interactions with nodes in the NFκB pathway

Besides the effects on cell movement and invasion, loss of function of this gene led to increased growth *in vivo* and *in vitro* (Fig. 1E and Fig. S2A). Transcriptomic analysis of *RBM10*-null compared to RBM10 re-expressing PE121410 cells showed enrichment in functional annotations for cell proliferation with concomitant inhibition of apoptotic and autophagic cancer cell death-related functions and of canonical cell cycle checkpoint-related pathways (Fig. 3A and Fig. S3A-B). To identify mediators of cell growth and anti-apoptotic effects caused by RBM10- loss we performed a genome-wide CRISPR/Cas9 dropout screen containing 77,441 single guide RNAs (sgRNAs) targeting 19,115 genes (Sanson et al., 2018). Empty vector (pLVX) and RBM10-expressing KTC1 cells were transduced with the sgRNA library at a low multiplicity of infection (MOIL=L0.3) followed by hygromycin selection and maintenance in culture for 20 days to enable changes in sgRNA abundance (Fig. 5A). We identified 987 genes significantly depleted in KTC1-pLVX vs KTC1-RBM10 expressing cells (p<0.05) (Table S2). Of note, sgRNAs targeting *RBM5*, a paralog of *RBM10* with which it shares overlapping RNA targets (Sun et al., 2017, LoiselleRoy and Sutherland, 2017), was among the top depleted hits suggesting that co-deletion of these two AS genes could lead to synthetic lethality. KEGG pathway analysis revealed that the depleted hits in KTC1-pLVX cells were associated with pathways involving NFκB activation, including the terms NFκB signaling pathway, RIG-1- like receptor signaling, mitophagy and viral infection-induced signaling (Fig. 5B).

**Figure 5:**
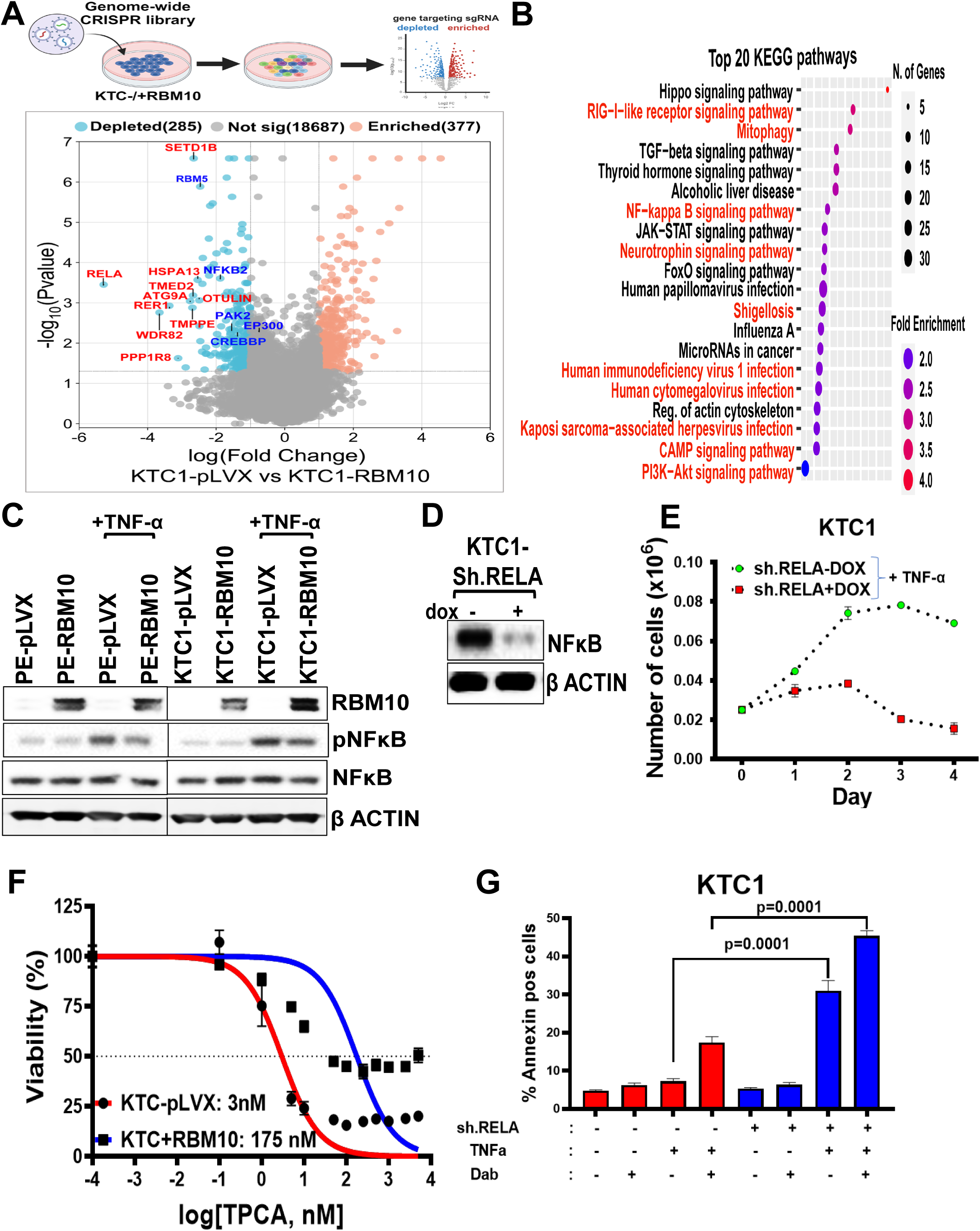
*Synthetic lethal interaction of RBM10 loss with NFκB:* (A) CRISPR/Cas9 dropout screen strategy to identify genes required for cell survival in *RBM10-null* KTC1 cells; Illustration by Biorender (*top*). Volcano plot (*bottom*) showing relative fold-change of depleted genes (light blue) in pLVX vs RBM10-expressing KTC1 cells calculated from an average of 4 sgRNAs targeting each gene. Gene label, red: Top 10 depleted genes; blue: other relevant hits. (B) KEGG pathway enrichment analysis performed in ShinyGO tool showing activation of NFκB and NFκB-related pathways (red). (C) Western blot of pNFκB (p65) and total NFκB in PE and KTC1 cells at baseline and in the presence of TNF-α. (D) Western blot demonstrating dox-induced KD of RELA (NFκB) in KTC1 cells. (E) Effect dox-induced KD of RELA on growth of *RBM10-*mutant KTC1 cells. (F) Dose-response curves of KTC1 cells -/+RBM10 treated with various concentrations of TPCA in the presence of TNFα in 1% FBS. (G) Effect of RELA KD on apoptosis in the presence or absence of TNFα (100ng/ml) and dabrafenib (200nM) as determined by annexin flow cytometry in KTC1 cells.

*RELA* (NFκB) topped the list of dropout hits within the NFκB signaling network in terms of negative fold-change in KTC1-pLVX vs KTC1-RBM10 cells (logFC=-5.28; p=0.00035) (Fig. 5A) and scored in all NFκB-relevant KEGG pathways (Table S3). In addition to *RELA*, *NFKB2*, which binds to RELB to mediate non-canonical NFκB signaling, was also among the significantly depleted hits. We were unable to identify signaling nodes upstream of RELA subject to RBM10-dependent cassette exon AS. Interestingly, CREBBP and its homolog p300, transcriptional coactivators and lysine acetyltransferases that play an important role in NFκB- mediated transcription, were among the dropout hits in the CRISPR screen. CREBBP exhibits AS of exon 11 in an RBM10-dependent manner (Fig. S5A). RELA competes with CREB for binding to the amino-terminal domain of CREBBP(Parry and Mackman, 1997) , but the domain encoded by exon 11 is distal to the interacting region, so the functional consequences of this exon inclusion are unknown.

The association of RBM10 loss with NFκB activation is supported by the fact that PE121410 and KTC1 cells show higher NFκB transcriptional output scores as compared to their respective isogenic RBM10-expressing controls (Fig. S5B). TNFα-induced phospho-p65 levels were attenuated by RBM10 re-expression in PE121410 and KTC1 cells, consistent with de-repression of NFκB signaling by RBM10 loss (Fig. 5C). Dox-induced knockdown of RELA decreased growth of Rbm10-null KTC1 cells (Fig. 5D-E), which also displayed higher sensitivity to the NFκB inhibitor TPCA-1 compared to KTC1-RBM10 cells (Fig. 5F). Furthermore, treatment with TNFα, alone or in combination with dabrafenib, modestly induced cell death in KTC1 cells as assessed by flow cytometry for annexin (Fig. 5G). By contrast, dox-induced knockdown of RELA showed a marked induction of apoptosis by TNFα itself, which was further accentuated by combined treatment with dabrafenib. Taken together, these data indicate that NFκB activation in RBM10-deficient cells protects cells from TNFα induced apoptosis, and that genetic and pharmacological targeting of this pathway results in a synthetic lethal interaction.

## DISCUSSION

Most cancers exhibit widespread RNA splicing changes compared to the untransformed cells from which they are derived (Kahles et al., 2018). The mechanisms accounting for mRNA mis-splicing include somatic gain- or loss-of-function mutations of genes encoding key splicing regulatory proteins, altered expression of splicing regulators and cis-acting mutations that modify splicing of genes affected by these mutations (Bradley and Anczuków, 2023) (Dvinge et al., 2016). The mutual exclusivity of splicing genetic defects within a given tumor type suggests that cancers cells rely on genome-wide dysfunctional splicing to determine their phenotype (Seiler et al., 2018). *RBM10* mutations were noted to be enriched in patients dying of non-anaplastic thyroid cancer (Ibrahimpasic et al., 2017). Subsequently, a pan-cancer study of genes associated with metastatic disease in 25,000 patients revealed a significant increase in *RBM10* mutations in PTC patients with distant metastases genotyped at our institution, where thyroid cancer profiling is primarily performed in patients requiring systemic therapies for advanced disease (Nguyen et al., 2022). Interestingly, no association was found between *RBM10* mutations and metastatic disease in any other cancer type in this study, including in patients with NSCLC, despite the higher prevalence of *RBM10* mutations in lung compared to thyroid cancers (Imielinski et al., 2012). Among other AS changes, RBM10 loss in NSCLC results in inclusion of exon 9 of NUMB, which destabilizes this NOTCH signaling repressor leading to activation of the NOTCH pathway and stimulation of cell growth (Bechara et al., 2013). Enrichment of the exon 9 inclusion isoform of NUMB was also present in RBM10-mutant thyroid cancer cell lines, but by contrast to lung this was not associated with increased NOTCH pathway transcriptional output. Moreover, no NOTCH pathway genes scored in the CRISPR/Cas9 dropout screen for growth dependencies conferred by RBM10 loss. Thus, though many RBM10 AS targets may be common between tumor types, their phenotypic consequences appear to be lineage dependent.

RNA sequencing of human RBM10 isogenic thyroid cancer cell lines revealed that 2-6% of alternatively spliced cassette exons are differentially spliced following RBM10 loss, confirming that *RBM10* acts as a repressor of AS, as opposed to perturbations of core spliceosome factors such as *SF3B1, SRSF2, and U2AF1*, which disrupt global splicing and lead to widespread retention of constitutive introns or AS of normally constitutive junctions (Dvinge and Bradley, 2015). Identification of *RBM10* AS targets involved in disease pathogenesis is challenging, since individual cassette inclusion isoforms that are retained after RBM10 loss may be normally expressed in sub-stoichiometric ratios, thus precluding simple phenotypic annotation. Several lines of evidence point to a key role of RBM10 in modulating splicing of genes involved in promoting metastatic disease: 1. GO analysis of global transcriptomes as well as transcripts subject to RBM10-dependent AS in isogenic human thyroid cancer cell lines showed enrichment of pathways involved in cell adhesion, integrin binding and structural components of the cytoskeleton. 2. RBM10-deficient cells display potent activation of the G protein RAC1, which transduces signals emanating from extracellular matrix interactions with integrins to promote formation of lamellipodia and cell motility (Minden et al., 1995). 3. Deletion of RBM10-regulated exon-inclusion isoforms of the extracellular matrix protein tenascin C, the cell surface glycoprotein CD44 and the cytoskeletal protein vinculin each individually decrease either cell motility, invasiveness or both in RBM10-deficient human thyroid cancer cells. 4. Combined deletion of all three isoforms from a lung metastatic mouse *Hras^G12V^/Rbm10^flox^*cell line inhibits RAC1-GTP and its downstream signaling. 5. Whereas deletion of individual exon inclusion isoforms from *Rbm10* mutant cells was insufficient to abrogate development of lung metastases following tail vein injection, dual or triple isoform knockdown markedly inhibited metastatic burden. 6. This points to cooperative effects of at least three protein variants controlled by RBM10-dependent AS in both human and mouse thyroid cancers, which collectively impact a common signaling program driving cell motility and invasiveness.

The congenital malformations of TARP syndrome in humans include micrognathia and limb defects, such as talipes equinovarus and, less frequently, syndactyly (Kumps et al., 2021, Johnston et al., 2010, Niceta et al., 2019). Interestingly, murine Rbm10 is expressed in mid-gestation embryos in the 1^st^ and 2^nd^ branchial arches and in limb buds. Expression of Rbm10 in the 1^st^ branchial arch, which gives rise to the mandible, is highest at E9.5 and E10.5 and declines thereafter(Johnston et al., 2010). Although the mechanisms accounting for the underdeveloped jaw and cleft palate of TARP syndrome children are unknown, the loss of the temporal sequence of Rbm10 expression in these developing structures may account for this. It is also plausible that it may be mediated in part through AS of genes involved in migration and tissue remodeling, such as those we identified in thyroid cancers with RBM10 loss.

Numerous studies have highlighted the importance of TNC in metastasis (Lowy and Oskarsson, 2015, Sun et al., 2018, Oskarsson et al., 2011). It has been studied in mechanistic detail in breast cancer, where TNC expression in breast cancer cells promotes their survival and outgrowth in the metastatic niche(Oskarsson et al., 2011). The TNC oligomer ranges in size between 180 and 300 kDa. Nine of its 17 fibronectin type III repeats, encoded by exons 10-16, are subject to AS, generating a large variety of isoforms that are differentially expressed in a lineage-specific manner during development and in various pathological contexts (Guttery et al., 2010). The large size of the TNC oligomer has hampered rigorous structure-function studies, which have relied primarily on recombinant TNC fragments. Consistent with our findings, isoforms that include exon 16 are enriched in invasive breast cancers (Guttery et al., 2010).

CD44 is a cell surface glycoprotein monomer involved in cell adhesion expressed in multiple cell types, which serves as a canonical receptor for hyaluronic acid but also interacts with other proteins, including osteopontin and matrix metalloproteases(Aruffo et al., 1990) (PrimeauxGowrikumar and Dhawan, 2022). It is subject to extensive AS, which alters domains involved in critical protein and cell-cell interactions(PrimeauxGowrikumar and Dhawan, 2022).

Colorectal cancer cell lines showing phenotypic plasticity between epithelial and mesenchymal states *in vitro* express RNA binding proteins downstream of the EMT master transcription factor Zeb1, including ESRP1 and several members of the RBM family, which regulate AS of CD44 in ways that favor metastatic competency(Xu et al., 2022). ESPR1 also regulates splicing of CD44 into a variant containing exon 8, which promotes metastatic colonization of breast cancers in lung(Yae et al., 2012). This aligns well with our data showing that RBM10 loss results in retention of exon 8-containing CD44 transcripts, which favor development of lung metastases. Taken together, this is consistent with a process directed by transcriptional drivers of EMT, which modulate expression of RNA binding proteins that in turn generate AS events that alter key components of the proteome to promote metastases.

Vinculin is a key player in focal adhesion-mediated regulation of cell behavior (Humphries et al., 2007). Its exon inclusion isoform MVCL is preferentially expressed in smooth and cardiac muscle cells (Belkin et al., 1988b, Belkin et al., 1988a), where it negatively impacts F-actin bundling by VCL and increases cell migration velocity (Lee et al., 2019). Germline mutations within the 68-aminoacid MVCL insert are associated with hypertrophic and dilated cardiomyopathy, highlighting its functional significance (Maeda et al., 1997, Olson et al., 2002). We show that RBM10 loss alters the VCL/MVCL ratio to induce cell motility, an isoform switch that had not been previously implicated in cancer pathogenesis.

Besides its effects on cell migration and invasiveness, constitutive RAC1 activation can also promote cell growth(Ma et al., 2009) . Activating mutations of *RAC1* are found in patients with ATC(Pozdeyev et al., 2018) and have been implicated in acquired resistance of BRAF-mutant PTC to MAPK inhibitors and in promoting anaplastic transformation(Bagheri-Yarmand et al., 2021). Despite this the whole genome CRISPR screen for genes required for viability of RBM10-null thyroid cancer cells did not identify effectors in the RAC1 signaling pathway among its top hits (PAK2, a downstream effector of RAC1, was a notable exception). Instead, RELA was the highest scoring hit as well as several other NFκB-related genes, and KEGG analysis of the dropout genes primarily identified pathways associated with NFκB activation. Nuclear localization of p65-RELA in PTC and ATC has been shown by several groups(Pacifico et al., 2004, Mitsiades et al., 2006). In addition, NFκB activation controls thyroid cell growth, migration and invasiveness in subsets of human thyroid cancer cell lines(Cormier et al., 2023). Our data shows that RBM10 loss renders thyroid cancer cells dependent on the NFκB pathway for viability. This provides a possible therapeutic strategy for these aggressive cancers, for instance by targeting TNFα, which is both an activator and an effector of the NFκB pathway, most likely in the context of combination therapies(Yu et al., 2020).

In summary, RBM10 loss in thyroid cancer is associated with disease progression and promotion of metastatic fitness through exon inclusion events in transcripts encoding proteins involved in cell motility and invasiveness, which result in constitutive activation of RAC1. The tumor suppressor role of RBM10 may thus exert its effects through restricting these EMT-related properties and by impairing growth inputs transduced through cancer cell autonomous NFκB signaling.

## MATERIALS AND METHODS

### Patient tumor samples

After IRB approval, a total of 785 PTC, high-grade follicular cell-derived thyroid cancer (HGFCTC) and anaplastic thyroid cancer samples from 724 unique patient were included in this study. The samples were collected from 1999 to 2023 and were sequenced at Memorial Sloan Kettering Cancer Center from 2014 to 2023. HGFCTC was defined by histological and/or immunohistochemical evidence of follicular cell differentiation and presence of tumor necrosis and/or ≥ 5 mitoses per 10 high-power fields (×400) (Hiltzik et al., 2006). All tumors were profiled using the Memorial Sloan Kettering Integrated Molecular Profiling of Actionable Cancer Targets (MSK-IMPACT) clinical sequencing assay, a hybridization capture-based, next-generation sequencing platform (Cheng et al., 2015). Patient demographics, tumor histology, metastatic status, metastatic sites, treatment, and outcomes were determined by retrospective review of patient charts. Altogether 737/785 samples were subjected to mutation analysis after filtering out multiple samples taken from the same patients. For mutation analysis patient samples were sorted for the following histotypes and for the presence of distant metastatic disease: PTC with metastases (PTC_wMet; n=321); PTC without metastases (PTC_woMet; n=56); HGFCTC (n=224; all with metastatic disease); ATC (n=136; all with metastatic disease). In addition, among the 496 samples from TCGA (Cancer Genome Atlas Research, 2014), 466 with known clinical characteristics were sorted for PTC with metastases (n=8) and PTC without metastases (n=458). These were pooled with the respective MSK PTC cohorts for *RBM10* mutation analysis.

Oncoprint and lollipop plots for *RBM10*-mutations on the sorted MSK cohort were generated using cBioPortal (Gao et al., 2013). The association of *RBM10* mutation with metastatic disease was obtained by contingency analyses using GraphPad Prism and 2-sided Fisher’s exact test for statistical significance. The thyroid cancer groups with (TC_wMet) and without metastases (TC_woMet) were obtained by combining TCGA PTC and the MSK cohorts of PTC and HGFCTC.

### Generation of thyroid-specific Rbm10 knockout mice

We crossed *Tpo-Cre (Kusakabe et al., 2004)*, *FR-Hras^G12V^*(Chen et al., 2009)*, Rbm10^flox^* (Wang et al., 2023) and *LSL-eYFP* mice (Jackson Laboratory; stock number 007903) to generate quadruple *Tpo-Cre/eYFP/Hras/Rbm10* (TEHR) transgenics, and the following control lines: *Tpo-Cre/eYFP* (TE or WT), *Tpo-Cre/eYFP/Hras* (TEH) and *Tpo-Cre/eYFP/Rbm10* (TER). These multi-transgenic mice result in Tpo-Cre driven thyroid-specific loss of *Rbm10,* endogenous expression levels of Hras^G12V^ (Het or Homo) and expression of YFP in thyroid follicular cells. Unless indicated Hras^G12V^ are homozygous in the TEH and TEHR mice; Rbm10 in the TER and TEHR mice are floxed either 1 or 2 alleles depending on the sex of the mice (female: Rbm10^fl/fl^, male: Rbm10^fl/y^), herein referred as Rbm10^flox^ irrespective of sex. DNA extractions from the toe clips were used for genotyping. Genotypes were determined by PCR using primers listed in Table S4. Mice were in mixed genetic backgrounds. Animal care and all procedures were approved by the Memorial Sloan Kettering Cancer Center (MSKCC) Institutional Animal Care and Use Committee (IACUC).

### Ultrasound imaging

Mice were anesthetized by inhalation of 1-5–2.5% isoflurane with 2% O_2_, the neck fur removed with defoliating agent and placed on the heated stage. An aqueous ultrasonic gel was applied over the neck and thyroid tumors were imaged with the VisualSonics Vevo™ 770 In Vivo High-Resolution Micro-Imaging System (VisualSonics Inc, Toronto, Ontario, Canada). Using the Vevo™ 770 scan module the entire thyroid bed was imaged with captures every 250 microns. Using the instrument’s software, the volume was calculated by manually tracing the margin of the tumor every 250 microns.

### Histology

Thyroid cancer histological characterization was performed at 6 months to 1 year-old mice or when recommended by the Research Animal Resource Center (RARC) veterinary staff because of tumor burden. Dissected mouse thyroids and lungs were fixed in 4% paraformaldehyde, embedded in paraffin, sectioned, and stained with hematoxylin and eosin (H&E) by the MSK Molecular Cytology Core Facility. Histologic diagnosis was performed by two thyroid pathologists (B. Xu and R.A Ghossein) blinded to mouse genotype. Slides were scanned with Pannoramic Flash 250 (3DHistech), and whole thyroid lobes or regions of interest were viewed using CaseViewer and exported as tiff images.

### Cell Lines

KTC1, PE121410, HTH83, SW1736 and 8305C human thyroid cancer cell lines were grown in RPMI medium and HEK293FT cells in DMEM, and all were supplemented with 10% of FBS and 1% penicillin/streptomycin/L-glutamine (PSG; Gemini; #400-110) and maintained at 37°C and 5% CO2 in a humidified atmosphere. Cell lines were previously genotyped by targeted cancer exome sequencing (MSK-IMPACT platform)(Landa et al., 2019), tested negative for Mycoplasma and were authenticated using short tandem repeat and single-nucleotide polymorphism analyses. Mouse thyroid tumor cell lines were derived as previously described(Saqcena et al., 2021). Briefly, the *TEHR* mouse metastatic cell line 64860M was generated from a lung metastasis that was dissected, minced in F12 medium and resuspended in 10 mL of digestion medium (minimum essential media containing 112 U/mL type I collagenase; Worthington, cat. #CLS-1), 1.2 U/mL dispase (Gibco; catalog no. 17105-041), penicillin (50 U/mL), and streptomycin (50 mg/mL). Cells were incubated at 37°C for 60 minutes with vigorous shaking, after which cells were spun down and resuspended in Coon’s modified F12 medium. The cell suspension was then sorted for YFP to eliminate non-thyrocytes and YFP^+^ cells plated in F12 medium supplemented with 5% FBS and 1% PSG. Cells were passaged at least five times and tested for Mycoplasma prior to use in experiments.

### RBM10 overexpression and knockdown

KTC1 and PE121410 cells with *RBM10* loss-of-function mutations were used to generate dox-inducible RBM10 expressing cells (KTC-RBM10 and PE-RBM10) by transducing the pLVX-Tet-On Advanced vector system (Clontech) with RBM10 cDNA (NM_005676, Origene) cloned into the pLVX-tight-Puro vector. Empty pLVX-puro vector-transduced cells were used as controls (KTC-pLVX and PE-pLVX). The *Hras^G12V^/Rbm10^flox^* mouse metastatic 64860M cells were used to generate Rbm10-expressing cells by transducing pLVX-puro vector cloned with mouse Rbm10 cDNA (NM_145627, Origene) (64860M-Rbm10). Control cells were transduced with empty vector (64860M-pLVX). RBM10 knockdown (KD) was performed in selected human thyroid cancer cells: Hth83, SW1736 and 8305C. We used MISSION® shRNA lentiviral vectors (pLKO_TRC005): TRCN0000233277 (sh.RBM10-A) and TRCN0000233278 (sh.RBM10-B) and the empty pLKO vector (sh.cont), all purchased from Sigma. To create cells with dox-inducible RELA shRNAs, we first transduced KTC1 cells with pLVX-Tet-On Advanced vector to obtain KTC1-rtTA cells, which were then infected with pLV-miR30, TRE-driven dual-shRNAs targeting human *RELA,* which were custom-designed and synthesized by VectorBuilder Inc. (Table S5). For lentiviral production, HEK293FT cells were transfected with lentiviral constructs using the Mission Lentiviral Packing Mix (Sigma). After media change at 24h viral supernatant was collected at 48h and 72h post-transfection, the lentiviral collections pooled, filtered (0.45 µm) and stored at −80°C. Stable lines were generated by infecting the target cells with the corresponding viral supernatants in the presence of 8μg/mL polybrene (Sigma) overnight. After 24h recovery in complete medium, cells were selected in 1μg/mL puromycin with or without 500 μg/mL G418, as required.

Expression of the target protein (RBM10 or RELA) was tested in multiple clones or in mass cultures. Depending on the construct, samples with> 80% KD (for RBM10 or RELA shRNAs) or near-endogenous levels of RBM10 expression (for dox-inducible RBM10) as determined by immunoblotting were selected for further experiments.

### Exon-inclusion isoform specific KD

Isoform-specific KDs were generated via targeted knockdown of the indicated exon inclusion isoforms in human RBM10-mutant KTC1 cells and in Luc^+^64860M cells. We designed shRNAmir hairpins with SplashRNA (http://splashrna.mskcc.org/) targeting the following human/mouse exons: human/mouse exon 19 inclusion transcripts of VCL (VCL_Ex19/Vcl_Ex19), exon 16 inclusion transcript of human TNC (TNC_Ex16) and the mouse equivalent exon 14 of Tnc (Tnc_Ex14), and human/mouse exon 8 inclusion transcripts of CD44. A similar approach was used to design 3 additional shRNAs against VCL_Ex19, CD44_Ex11, CD44_EX15 and TNC_Ex7. The 97-bp shRNAmirs were synthesized, PCR-amplified and cloned into the lentiviral miR-E based SREP (pRRL) vector as previously described(Fellmann et al., 2013).

For combined isoform knockdown in Luc^+^64860M mouse cells, we used one of the above validated shRNAmirs for each target: Vcl_Ex19, Cd44_Ex8 and Tnc_Ex14, which were custom designed into lentiviral miR30-based dual and triple shRNA expression vectors (pLV[miR30]- Puro-SFFV>DsRed_shRNAs) constructed by VectorBuilder. The following dual- and triple- isoform specific shRNA vectors targeting the transcripts expressing the following mouse gene exons were generated: shVcl_Ex19/shCd44_Ex8, shVcl_Ex19/shTnc_Ex14, shCd44_Ex8/shTnc_Ex14 and shVcl_Ex19/shCd44_Ex8/shTnc_Ex14 (sh.3KD). Sh.Scramble (sh.Scr) was used as control vector. All shRNAs and additional vector information are described in Table S5. Human and mouse cells with stable isoform-specific KDs were created via transduction of the corresponding lentiviral particles. Cells were sorted for RFP as a reporter of shRNA expression to increase KD efficiency.

### Western blotting

Cells were lysed in 1 x RIPA buffer (Millipore) supplemented with protease (Roche) and phosphatase inhibitor cocktails I and II (Sigma). Mouse thyroid tumors were homogenized in 1 x Lysis Buffer (containing 10 mmol Tris-HCl, 5 mmol EDTA, 4 mmol EGTA and 1% Triton-X100) with protease/phosphatase inhibitors. Lysates were briefly sonicated to disrupt the tissue and cleared by centrifugation. Protein concentrations were estimated by BCA kit (Thermo Scientific) on a microplate reader (SpectraMax M5); comparable amounts of proteins were subjected to SDS-PAGE using NuPAGE 4%–12% Bis–Tris gradient gels (Invitrogen) and transferred to PVDF membranes. Following overnight primary antibody, membranes were incubated with secondary antibodies coupled to horseradish peroxidase (HRP) or IRDye fluorophores for 1 h at room temperature. HRP probed blots were developed using enhanced chemiluminescence reagent (Amersham Biosciences), and signal was captured using the iBright CL1000 Imaging system (ThermoFisher Scientific). IRDye-probed blots were imaged using the LI-COR Odyssey imaging system (LI-COR Biosciences).

### Antibodies and other reagents

The following primary antibodies were used for Western blots at 1:1000 dilution, except where indicated. RBM10 (HPA034972) and β-actin (A2228; 1: 10,000) from Sigma-Aldrich. VCL(#13901), PAK1,2,3 (#2604), pPAK1-(Thr423)/PAK2-Thr402 (#2601), AKT (#2920), pAKT-Ser473 (#4051), NF-κB/p65(#8242) and pNFKB-S536 (#3033) from Cell Signaling Technology. The secondary antibodies were used at 1: 5,000 dilutions. We used the following HRP-conjugated antibodies: goat anti-rabbit (Santa Cruz; sc-2004) and goat anti-mouse (Santa Cruz; sc-2031) and the following IRDye fluorophore-conjugated antibodies: IRDye 800CW Goat anti-Rabbit IgG (LI-COR; 926-32211), IRDye 800CW Goat anti-Mouse IgG (LI-COR; 926-32210), IRDye 680RD Goat anti-Rabbit IgG (LI-COR; 926-68071), and IRDye 680RD Goat anti-Mouse IgG (LI-COR; 926-68070). We also used the following additional reagents *in vitro*: doxycycline (2 μg/mL) from Sigma, TPCA-1 (#S2824) from Selleckchem, recombinant human TNFα (#16769) from Cell Signaling.

### RAC1-GTP pulldown assay

RAC1 activation was determined using the active Rac1 pull-down and detection Kit (ThermoFisher Scientific). Briefly cells were seeded at 40% confluence, treated with doxycycline where needed, and serum starved (1% FBS) for at least 48h prior to collection. Cells were then washed with ice-cold PBS, lysed on ice, and at least 500µg lysate was used for RAC1-GTP pull down following the manufacturer’s protocol. The eluted RAC1- GTP was denatured, electrophoresed in SDS-PAGE and Western blotted. Western blots of input lysate were probed for total RAC1 for normalization, total and pPAK (pPAK1- (Thr423)/PAK2-Thr402) and total and pAKT-S473 to assess RAC1 downstream signaling.

### Reverse transcriptase PCR (RT-PCR)

Total RNA from isogenic cell lines was extracted using the RNeasy mini kit (Qiagen). Comparable amounts of RNA (500ng-1 μg) were subjected to DNase I (Invitrogen) treatment and reverse transcribed using SuperScript III Reverse Transcriptase (Invitrogen) following the manufacturer’s protocol. cDNA was diluted at 1:15 and 2ul used as a template for either RT-PCR or qRT-PCR reactions. RT-PCR was performed using REDTaq® ReadyMix™ (Sigma) on a thermocycler (Eppendorf) and qualitative assessment of PCR products were performed by resolving the amplicons in 3% agarose gel. For quantitative assessment, qRT-PCR was performed using the Power SYBR Green PCR Master Mix (Applied Biosystems) on QuantStudio 7 pro (Applied Biosystems). For gene expression quantifications the Ct values of the target genes were normalized to GAPDH (human) or Hprt (mouse), and for exon inclusion isoform quantifications the relative ratios were calculated by normalization to corresponding constitutive exons as previously described(Wang et al., 2013). The PCR primers are listed in Table S4.

### Cell migration and invasion assay

Cell migration was assayed by time-lapse imaging and tracking of single cells over time as previously described(Lee et al., 2019). Briefly 50,000 cells/well were seeded in a 6-well culture dish overnight prior to imaging. Cells were imaged with a 10x/0.45NA objective on a Zeiss Axio Observer.Z1 microscope using the imaging software ZEN Blue 2.3 Pro for 16 h with 8-minute intervals at 25 random positions in each well. Cells were maintained at 37°C with 5% CO2 during the imaging period. Single cell tracking was performed using the TrackMate plugin in FIJI, in which single cells were tracked based on displacement of each object over time. Cells that experience cell division, cell death, a collision event, or migrated out of the field of view were excluded. To compute cell velocity, raw tracking data was analyzed using MATLAB. Time-lapse imaging was performed by the MSK Molecular Cytology Core Facility.

For invasion assays, cells were trypsinized, washed in PBS, resuspended in 0.5% FBS and 25,000 cells (0.5Lml) plated in the top compartment of an 8.0 µm fluorescence blocking PET membrane of the Corning® BioCoat™ Matrigel® Invasion Chambers in technical triplicates. Cells that invaded into the bottom membrane were imaged and counted after 24h using a fluorescence microscope.

### In vivo metastasis assays

Metastases experiments were conducted in 6-8 week-old female immunocompromised Rag1 deficient mice (strain: B6.129S7-Rag1tm1Mom/J; Jackson laboratories) in accordance with a protocol approved by the MSKCC Institutional Animal Care and Use Committee. To assess metastatic fitness, we orthotopically implanted luciferase-transduced 64860M-pLVX and 64860M-Rbm10 cells into a thyroid lobe. Mice were anesthetized by inhalation of 1-5–2.5% isoflurane with 2% O2, neck hair removed with defoliating agent and placed on the heated stage of the ultrasound. The site of injection was cleansed with 70% ethanol-soaked gauze. An aqueous ultrasonic gel was applied over the neck and thyroid lobes were visualized by the ultrasound. By ultrasound guidance, 5μl of tumor cell suspension in PBS containing 10^4^ cells were injected into a thyroid lobe with a Hamilton syringe loaded with a 30-gauge sterile needle. The primary orthotopic tumors were monitored by weekly IVIS imaging, and the lung metastases by H&E staining of the lungs following sacrifice. Tumor cell tail vein (TV) injections were performed to assess the relative lung metastatic efficiency of 64860M cells engineered to re-express Rbm10 and shRNAmir-mediated KDs of individual or combined exon inclusion isoforms (Vcl_Ex19, Cd44_Ex8, Tnc_Ex14). Prior to IV injections mice were warmed under a heat lamp for 10 min, and 500,000 cells resuspended in 200ul PBS injected via the lateral tail vein of 6–8-weeks-old mice. Orthotopic and tail-vein implanted mouse lung metastases were assessed by serial IVIS imaging after administration of 2mg/mouse D-Luciferin (GOLDBIO). Bioluminescence images were captured using the IVIS spectrum CT In Vivo imaging system (Caliper Life Sciences), and quantified using Living Image software, version 2.60 by monitoring the total flux (photon/sec) in the tumor region of interest (ROI).

### RNAseq and splicing analysis

RNA was extracted from the following isogenic RBM10 overexpressing or KD cell lines using the Qiagen RNeasy extraction kit: KTC1, KTC1-RBM10, PE121410, PE121410-RBM10, 8305C-sh.Ctrl, 8305C-sh.RBM10, SW1736-sh.Ctrl, SW1736-sh.RBM10 and HTH83-sh.Ctrl, HTH83-sh.RBM10. After RiboGreen quantification and quality control by Agilent Fragment Analyzer, 500 ng of total RNA with RQN values of 9.1-10 underwent polyA selection and TruSeq library preparation according to instructions provided by Illumina (TruSeq Stranded mRNA LT Kit, catalog # RS-122-2102), with 8 cycles of PCR. Samples were barcoded and run on a HiSeq in a PE125 run, using the HiSeq 3000/4000 SBS Kit (Illumina). An average of 100 million paired reads was generated per sample. Ribosomal reads represented 1.3-8.4% of the total reads and the percent of mRNA bases averaged 73%.

Differential splicing analysis was performed as previously described(Dvinge et al., 2014). In brief, RNA-seq reads were aligned to the hg19/GRCh37 genome assembly using a annotation that consisted of a combination of annotations from UCSC knownGene(Meyer et al., 2013), Ensembl(Flicek et al., 2013), and MISO isoforms(Katz et al., 2010). Reads were aligned with RSEM(Li and Dewey, 2011), Bowtie(Langmead et al., 2009), and TopHat(TrapnellPachter and Salzberg, 2009). PSI values were then computed using MISO v.2.0(Katz et al., 2010).

Additionally, alternate splicing analysis and differential gene expression in the RNAseq of RBM10 isogenic KTC1 and PE121410 cells were performed using Partek Flow version 6.0.17.0614 (www.partek.com). Briefly, raw reads were aligned to human assembly hg19 using STAR aligner (STAR-2.5.3a) with default parameters. The aligned reads were quantified to Partek E/M annotation model using hg19 assembly generating gene and transcript counts. The transcript counts were subject to “Detect alt-splicing”, an ANOVA based Partek algorithm to detect genes with multiple isoforms and determine their expression changes. Detect alt-splicing with default parameters was applied on the following comparators by grouping RBM10-mutant vs the corresponding RBM10 re-expressing counterpart. Differentially expressed alternatively spliced transcripts (Table S1) were visualized by a volcano plot. Gene Ontology analysis on RBM10 induced differentially spliced targets were performed using GOrilla(Eden et al., 2009).

For differential gene expression quantification the Deseq2 algorithm was used for the comparators KTC1 vs KTC1-RBM10 and PE121410 vs PE1214410-RBM10. The differentially expressed gene list was used to run the Ingenuity Pathway Analysis (Qiagen). A pathway enrichment plot was constructed using the SRplot online tool(Tang et al., 2023).

### Whole exome CRISPR Screen

Prior to the screen, KTC1-pLVX and KTC1-RBM10 cells were transduced with Cas9. Cas9-expressing cells were then infected with the Brunello sgRNA library that consisting of 4 distinct sgRNAs/gene using a low multiplicity of infection (MOI) (∼0.3) to obtain a single sgRNA per cell. For MOI determination we transduced target cells with various volumes of the lentiviral library supernatant to obtain 30% infectivity based on survival following antibiotic selection. On day-5 post-transduction ∼75 million cells per replicate (x 3) were harvested with the goal of obtaining 1000 cells (1000X) expressing each sgRNA and the cell pellet frozen to assess the baseline sgRNA coverage. The remaining cells were serially passaged to maintain 750-1000X cells/sgRNA until day 24 post-transduction, when ∼75 million cells/replicate were harvested to determine the end point sgRNA distribution. Cell pellets were lysed, and genomic DNA extracted (Qiagen) and quantified by Qubit (ThermoScientific). A quantity of gDNA covering 1000X representation of sgRNAs was PCR amplified to add Illumina adapters and multiplexing barcodes. Amplicons were quantified by Qubit and Bioanalyzer (Agilent) and sequenced on Illumina HiSeq 2500. FASTQ pre-processing and guide abundance was determined using the MAGeCK count command. The 5’ trim length was automatically detected by MAGeCK, and a normalized count file was generated with median normalization. Sample QC showed >85% mapped reads and a minimum sample correlation of 0.8 for all replicates. Enrichment was determined using the MAGeCK RRA (robust rank aggregation) function to obtain gene-level enrichment scores, and p-values were determined by permutation. All analyses were performed using MAGeCK v.0.5.9. The differential sgRNA enrichments visualized by volcano plot using the SRplot online tool. KEGG pathway enrichment analysis was performed using the ShinyGO 0.77(GeJung and Yao, 2020). The 987 depleted sgRNAs in KTC1-pLVX vs KTC1-RBM10 with p values <0.05 (Table S2 were used as input to nominate the top 20 pathways with FDR < 0.2. A pathway enrichment plot sorted by fold-enrichment was constructed using the SRplot online tool.

### IC50 measurements

Cells plated in 96-well plates were exposed to the IKK2 inhibitor TPCA-1 at various concentrations with at least four technical replicates per concentration. On day 5 post-treatment, CellTiter Glo reagent (Promega) was used to determine cell viability as per the manufacturer’s protocol. Absolute viability values were converted to percentage viability compared to DMSO treatment. IC50 curves and values were generated on GraphPad Prism V8.0 using non-linear fit of log (inhibitor) vs response (three parameters).

### Annexin V assay

Apoptosis induced by dox-inducible RELA KD in KTC1 cells was determined using Annexin V-APC (BD Bioscience) and Annexin-binding Buffer (Invitrogen) according to the manufacturer’s specifications. KTC1-shRELA cells were plated with and without doxycycline in the indicated conditions. Cells harvested after 36h were subjected to annexin staining and captured by flow cytometry, and data analyzed using FlowJo software.

### Statistical Analyses

Statistical analysis for the CRISPR whole exome screen and RNAseq experiments is described in their respective sections. Statistical analysis of *in vitro* and *in vivo* studies was performed using two-tailed non-parametric t-tests with GraphPad Prism version 7.0. Significance was established as p<0.05.

### Online supplemental materials

Figure S1 shows molecular features associated with RBM10 mutations in human patients and histological characteristics of *Hras^G12V^/Rbm10^KO^* mouse thyroid cancer. Figure S2 demonstrates effects of RBM10-loss on thyroid cancer cell growth *in vitro* and lists AS targets of RBM10. Figure S3 shows inhibition of cell death-related pathways by RBM10 loss in isogenic PE121410 cells by Ingenuity Pathway Analysis. Figure S4 demonstrates metastases rescue by Rbm10 re-expression and validation of isoform specific KDs in 64860M cells. Figure S5 shows RBM10 effects on AS of CREBBP and NFκB transcriptional output. Table S1 lists AS transcripts differentially expressed in RBM10-null cells (KTC1 and PE121410) compared to their re-expression counterparts using Partek Alt-splicing analysis. Table S2 lists the drop-out genes in KTC1-pLVX Vs KTC1-RBM10 cells from the whole genome CRISPR screen. Table S3 shows KEGG pathways associated with CRISPR- Cas9 dropout hits, highlighting the involvement of the *NFKB* gene (*RELA*) (red) and CREBBP (blue) within the top 20 KEGG pathways enriched in the KTC1-cell CRISPR screen. Table S4 lists the primers used for genotyping, RT-PCR and qRT-PCR. Table S5 has the plasmid vector information used in the study.

## ACKNOWLEDGEMENTS

Supported by R01 CA50706-31, R01 CA255211-03 and R01 CA249663-03 (to JAF) and Cancer Center Support Grant P30 CA008748-58 (Selwyn Vickers). A.R.G wishes to thank the 2016 Royal Australasian College of Surgeons Foundation for Surgery Tour de Cure Cancer Research Scholarship and National Health and Medical Research Council Neil Hamilton Fairley Early Career Fellowship (RG183080) for salary support for this project. We are grateful for the contributions of the RNAi Core facility for the CRISPR screen, the Molecular Cytology, Small Animal Imaging, Anti-tumor Assessment and the Integrated Genomics Operation Core Facilities, funded by the NCI Cancer Center Support Grant (CCSG, P30 CA08748), Cycle for Survival, and the Marie-Josée and Henry R. Kravis Center for Molecular Oncology.

## AUTHOR CONTRIBUTIONS

G.P.K. and J.A.F. designed the study. G.P.K., A.G., N.S.-C., D.V., V.T. and A.A.-R. performed *in vitro* experiments. G.P.K., A.G., B.R.U., N.S.-C., D.V., K.B., P.P.T. and E.d.S. performed *in vivo* experiments. O.A.-W and J.A.F generated the Rbm10 GEM model. B.X. and R.A.G. performed histologic diagnosis of mouse tumors. Y.L. and L.B. identified and analyzed patient data. G.P.K. and R.K.B. performed computational analyses. G.P.K., A.G., M.S., J.A.K. and J.A.F. were involved in experimental design and data interpretation. G.P.K. and J.A.F. wrote the manuscript with input from all authors.

