## Supplementary material for "RBM10 loss induces aberrant splicing of cytoskeletal and extracellular matrix mRNAs and promotes metastatic fitness": Figure S1

### Figure S1: Molecular features associated with RBM10 mutations.

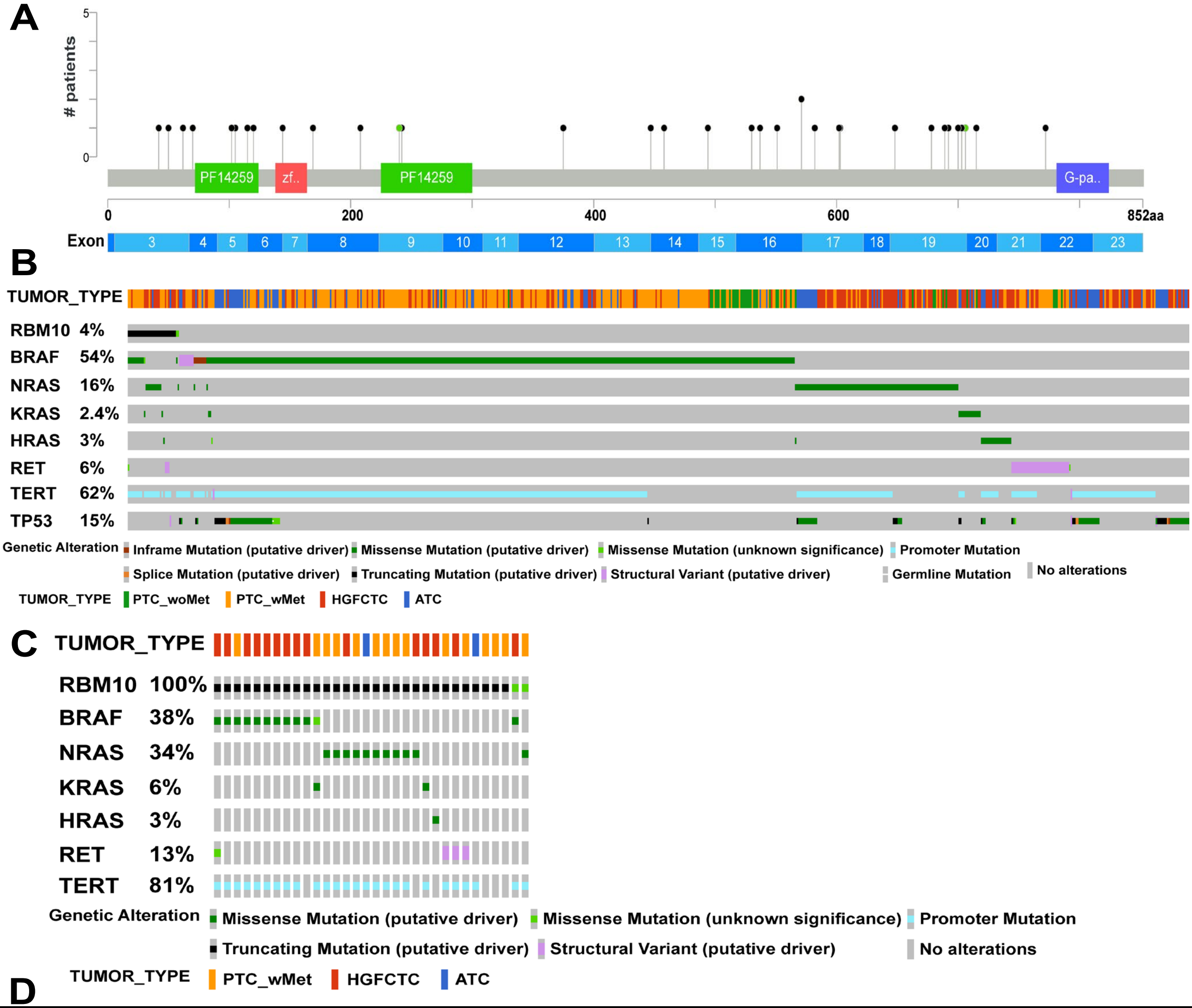


D

| | 6-8 months | | | | | |
|---|---|---|---|---|---|---|
| Genotype | Normal thyroid | PTC early - n (%) | PTC-n (%) | ATC-n (%) | Metastases | Total - n |
| *TEH* | nd | nd | nd | nd | nd | nd |
| *TER* | nd | nd | nd | nd | nd | nd |
| *TEHR* | 4 (11%) | 9 (26%) | 14 (40%) | 8 (23%) | 0 | 35 |

| | 9-12 months | | | | | |
|---|---|---|---|---|---|---|
| Genotype | Normal thyroid | PTC early - n (%) | PTC-n (%) | ATC-n (%) | Metastases | Total - n |
| *TEH* | 13 (100%) | 0 | 0 | 0 | 0 | 13 |
| *TER* | 26 (100%) | 0 | 0 | 0 | 0 | 26 |
| *TEHR* | 1 (3%) | 3 (9%) | 4 (12%) | 26 (76%) | 6 (18%) | 34 |

**Figure S1:** (A) Lollipop plot showing distribution and type of *RBM10* mutations. Black: nonsense; Green: missense mutations. (B) Oncoprint showing mutation frequency of *RBM10* and the indicated drivers in the MSK-clinical TC cohort (C) Oncoprint demonstrating co-occurrence of *RBM10* mutations with MAP kinase pathway alterations. (D) Thyroid cancer histological characteristics in mice with the indicated genotypes over time; nd - not determined
