## Supplementary material for "RBM10 loss induces aberrant splicing of cytoskeletal and extracellular matrix mRNAs and promotes metastatic fitness": Figure S2

### Figure S2: RBM10-loss confer growth advantage *in vitro.* AS targets of RBM10.

**A**

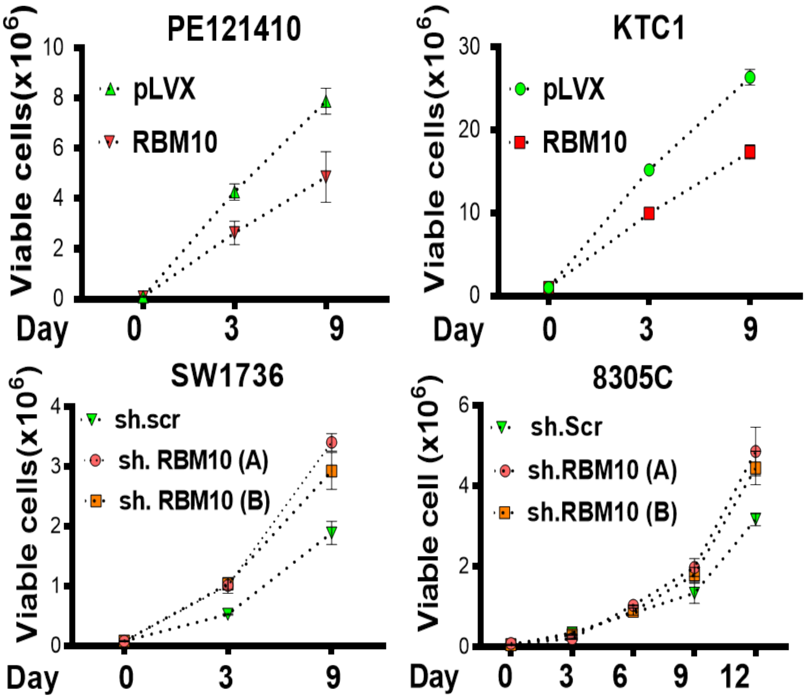


**B**

| | | | | | | | | | | | |
|---|---|---|---|---|---|---|---|---|---|---|---|
| AAMDC | ANKRD10 | BANP | CTDP1 | GGCT | MAP3K6 | NME2 | PLEKHH3 | RAPGEF3 | SUGP2 | TRPT1 | ZFAS1 |
| AC004166.7 | AP1B1 | BCL2L12 | DNM1 | GLE1 | MAP4K4 | NPIP | PLEKHM2 | RPLP1 | SUPT5H | UAP1 | ZNF33A |
| AC004381.6 | AP2M1 | C16orf13 | DNM2 | GOLGA2 | METTL23 | NSMF | PLSCR3 | SCRIB | SYNRG | UBAP2L | |
| AC069513.3 | APLP2 | CARM1 | DROSHA | HN1L | MORF4L2 | NUP62 | PMS2P5 | SEC22C | TBRG1 | UBXN11 | |
| AC103965.1 | APTX | CD46 | EIF4H | IKBKG | MPV17 | PAK4 | POLB | SEC31A | TMEM11 | UFD1L | |
| ACAD10 | ARAP1 | CDC16 | ERI3 | INF2 | MROH1 | PCBP2 | POLDIP3 | SLC25A19 | TMEM126B | UPP1 | |
| ACOT9 | ATG4B | CEP164 | EXOC7 | IP6K2 | MTMR3 | PEX5 | PRMT2 | SMPD4 | TMEM219 | USP21 | |
| ALAS1 | ATG9A | CLK2 | FAM189B | ISOC2 | MYO1B | PHLDB1 | PTK2 | SMUG1 | TNC | VKORC1 | |
| ALG2 | ATP5SL | CREBBP | FBXW11 | LETMD1 | NCAPG2 | PIGQ | PUF60 | STRADA | TPD52L2 | WNK1 | |
| ANAPC11 | BAG6 | CRYBB2P1 | FIZ1 | LRRC23 | NEIL2 | PKD1P1 | PVR | STYXL1 | TRIM65 | ZDHHC16 | |

**Figure S2:** (A) *Top:* Cell proliferation in *RBM10*-mutant thyroid cancer cells following expression of RBM10. *Bottom:* Effects of RBM10 silencing on growth of RBM10 wild-type thyroid cancer cell lines. (B) RBM10-dependent alternatively spliced (AS) genes common to 5 isogenic +/- RBM10 thyroid cancer cells.

### Figure S2: AS targets of RBM10.

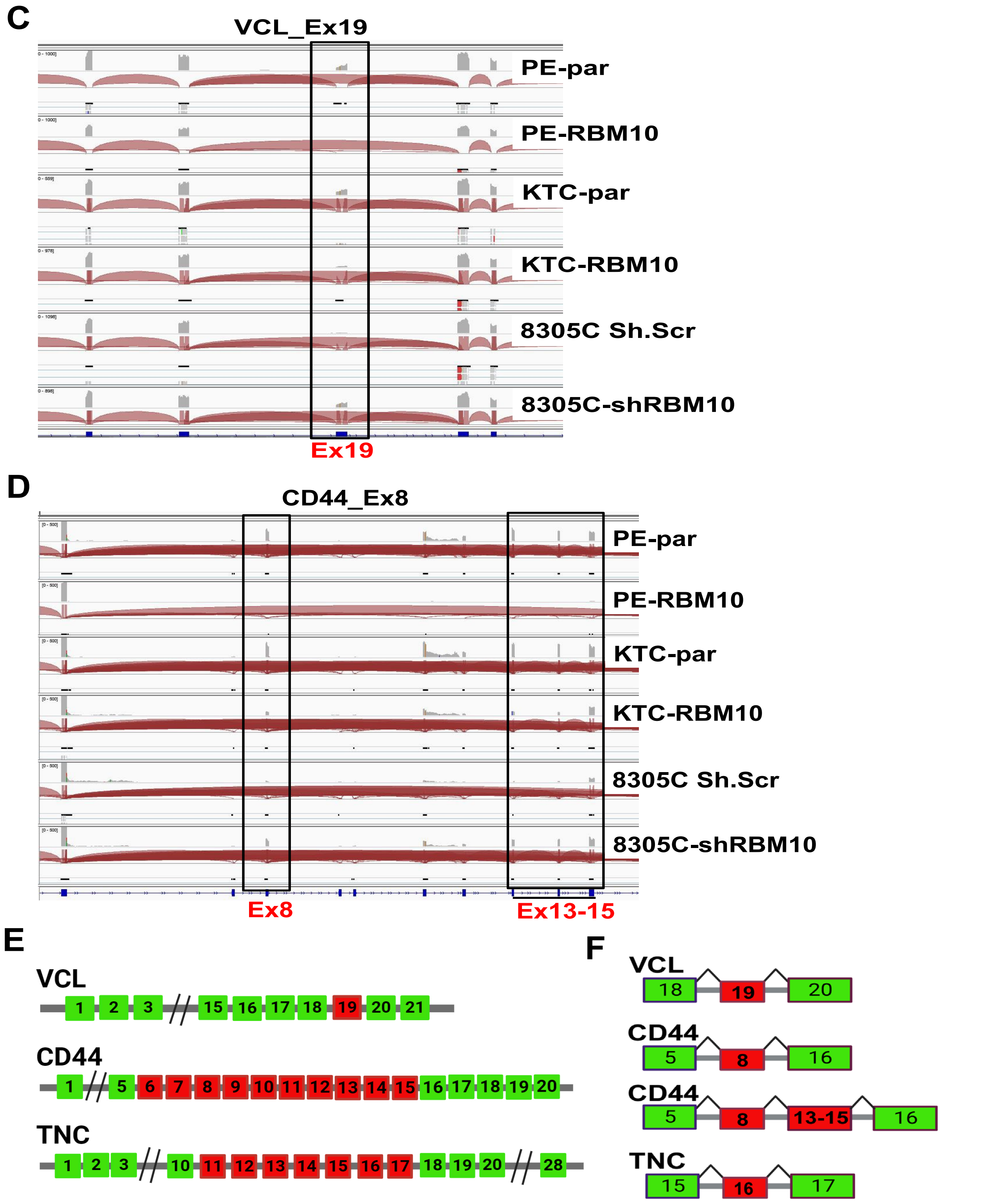


**Figure S2:** (C,D) IGV view of exon junctions in the RNAseq data of the indicated cells showing inclusion of exon 19 of vinculin (C) and exons 8 +13-15 of CD44 genes (D) in RBM10 mutant PE121410 and KTC1 cells and their exclusion following RBM10 expression, with reciprocal changes in RBM10-KD 8305C cells. (E) Scheme showing exon structure of *VCL*, *CD44* and *TNC* genes. (F) RBM10-targeted exon inclusion (red) isoforms of VCL, CD44 and TNC spliced-in with adjacent constitutive exons (green). PE par: parental PE121410; KTC par: KTC1 parental.
