## Supplementary material for "RBM10 loss induces aberrant splicing of cytoskeletal and extracellular matrix mRNAs and promotes metastatic fitness": Figure S3

### Figure S3: RNAseq analysis shows RBM10-loss induces deregulation of cell death and cell cycle pathways.

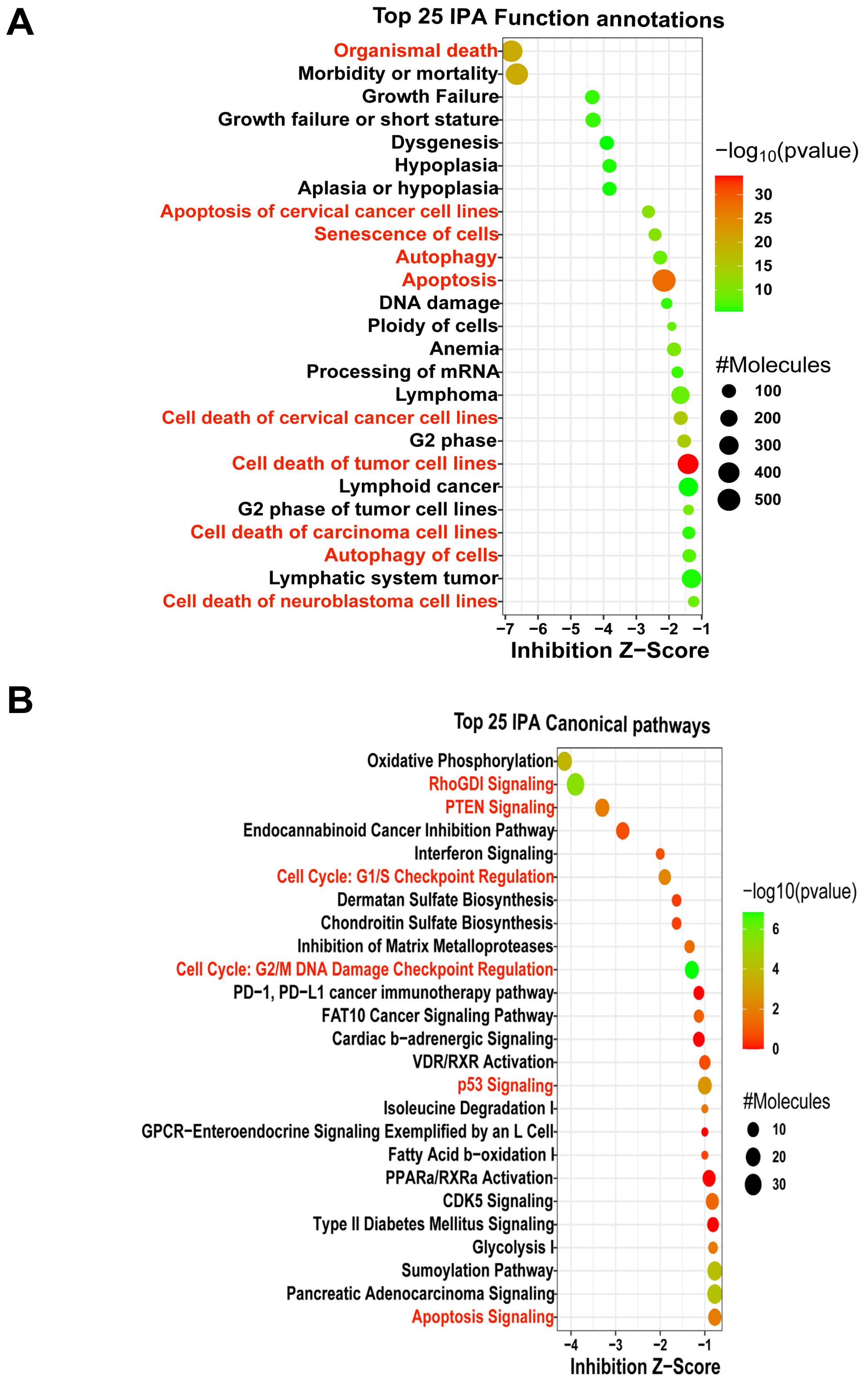


**Figure S3**: (A) IPA of RNAseq of PE121410 vs PE121410-RBM10 cells. (A) Inhibition of cellular functions associated with apoptotic and autophagic cancer cell death (red). (B) Enriched canonical pathways associated with inhibition of apoptosis, cell cycle checkpoints, and RhoGDI, an inhibitor of GDP dissociation in Rho family GTPases (red).
