## Supplementary material for "RBM10 loss induces aberrant splicing of cytoskeletal and extracellular matrix mRNAs and promotes metastatic fitness": Figure S4

### Figure S4: Suppression of metastases by RBM10 expression; Validation of isoform specific KDs.

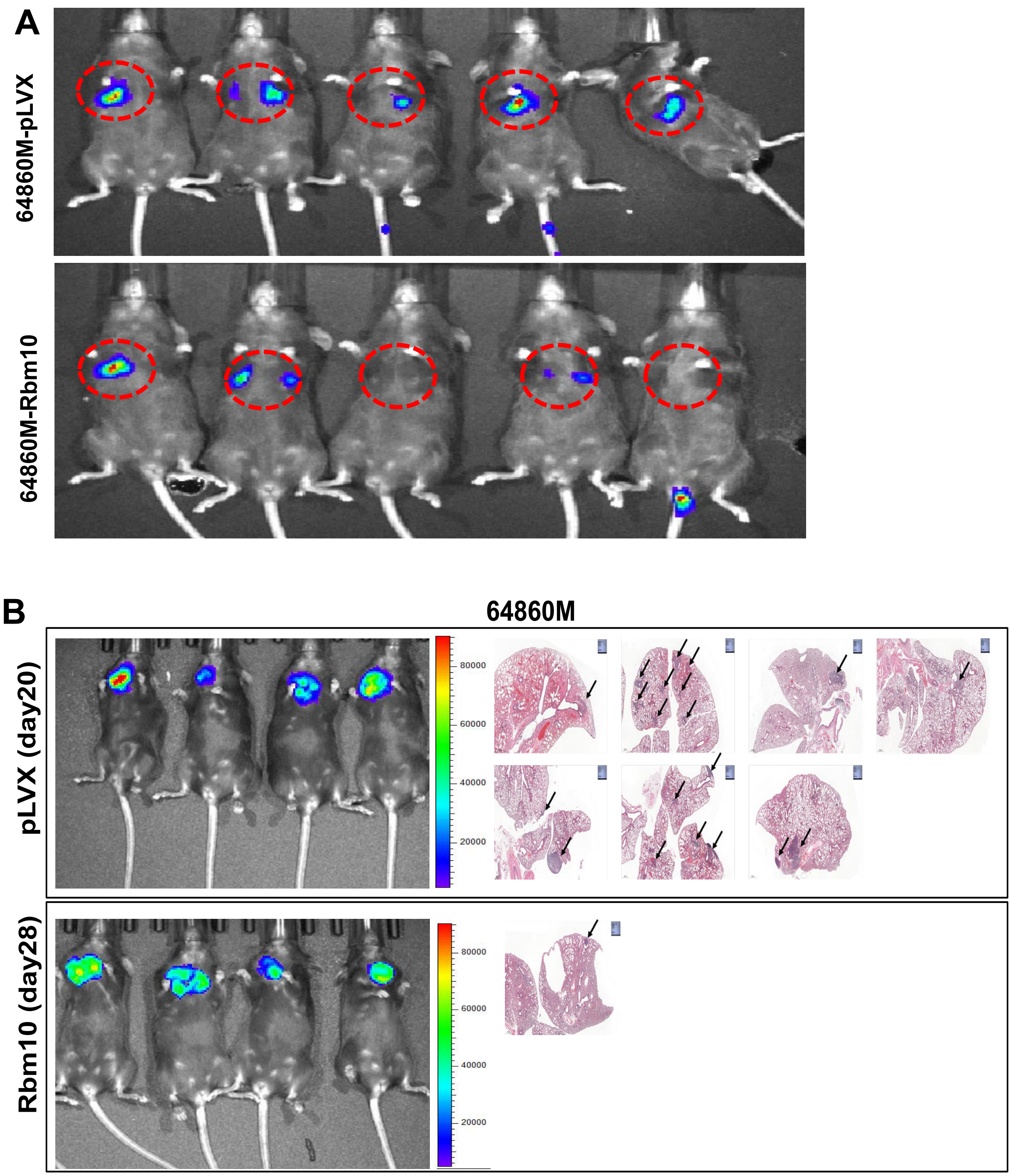


**Figure S4:** (A) Bioluminescence imaging of Luc+ 64860M cells 2 weeks after tail vein injection showing decreased lung colonization upon Rbm10 expression. (B) *Left:* Bioluminescence imaging of Luc+ 64860M cells 3-4 weeks after thyroid orthotopic implantation of 64860M cells -/+ Rbm10. Mice injected with Rbm10- expressing cells were sacrificed 8 days later to match tumor size with parental controls. *Right:* H&E stains of corresponding lung sections for each cohort. Arrows point to sites of lung metastatic colonization.

### Figure S4: Suppression of metastases by RBM10 expression; Validation of isoform specific KDs.

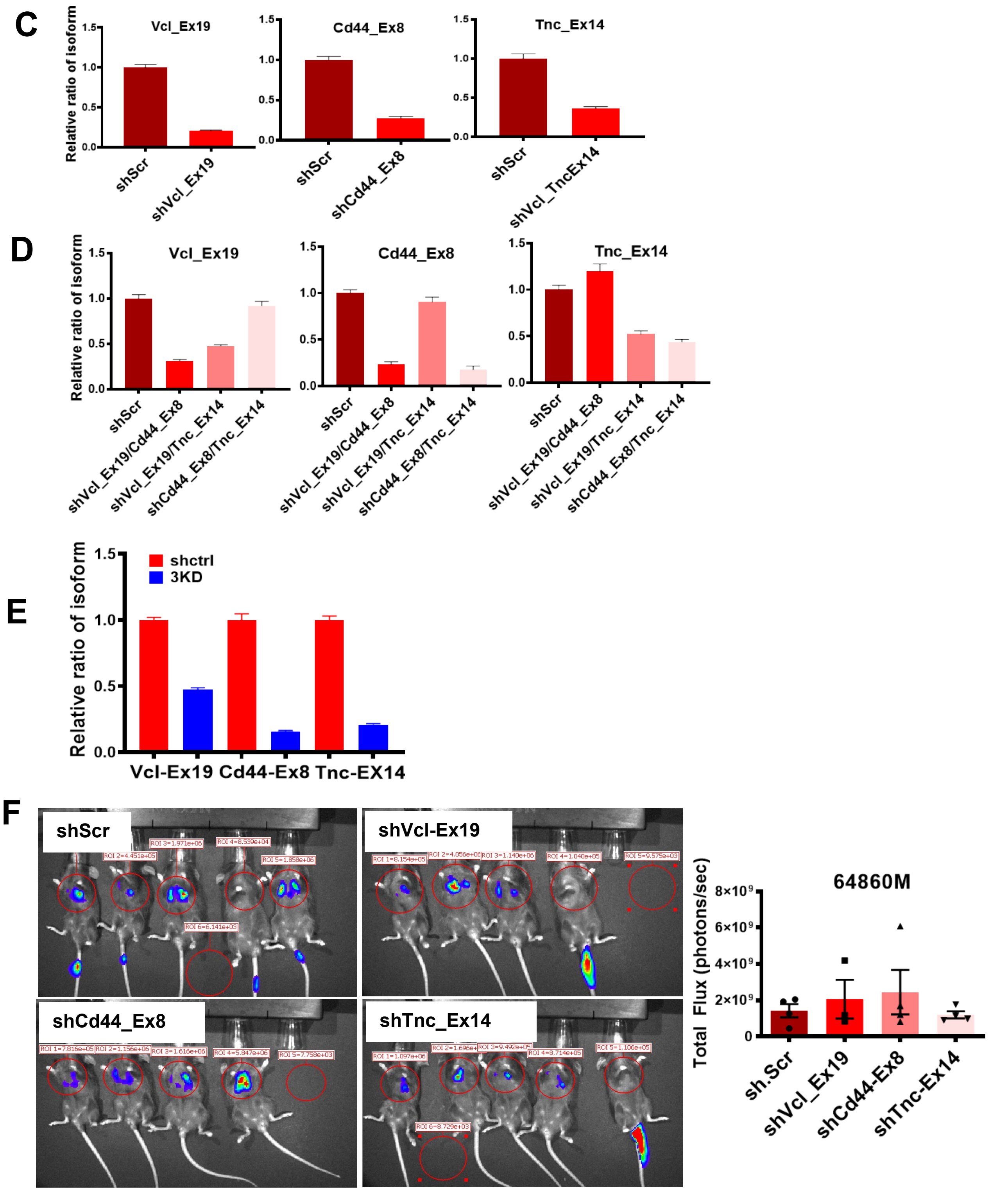


**Figure S4:** (C-E) Validation of isoform-specific shRNA KDs by qRT-PCR using junction-spanning primers of the indicated exons in 64860M cells; (C) single (D) dual and (E) triple isoform KD in 64860M vs sh-Scramble. (F) Bioluminescence imaging and quantification 2 weeks after TV injection of 64860M cells showing no changes in metastatic competence between sh.Scr and any of the individual isoform KDs.
