## Supplementary material for "RBM10 loss induces aberrant splicing of cytoskeletal and extracellular matrix mRNAs and promotes metastatic fitness": Figure S5

### Figure S5: RBM10 dependent CREBBP AS; RBM10 effects on NFκB transcriptional output.

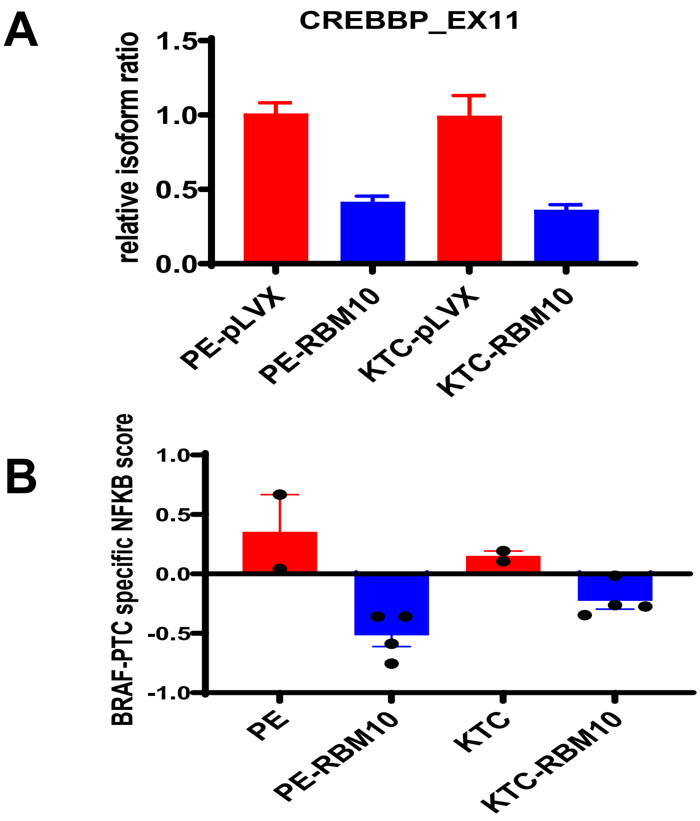


<u>**Figure S5:**</u> (A) Relative ratio of CREBBP exon 11 inclusion isoform determined by qRT-PCR in PE121410 and KTC1 cells -/+ RBM10 (B) NFκB activation score derived using a thyroid cancer-specific 50-gene signature (PMID: 37973791 ) in RBM10-null PE121410 and KTC1 cells vs their corresponding RBM10-expressing counterparts.
